# Non-canonical cleavage expands prime editing in 5’ direction

**DOI:** 10.1101/2025.05.20.655067

**Authors:** Pei-Ru Chen, Xian-Zheng Yuan, Shu-Guang Wang, Peng-Fei Xia

## Abstract

Cleavage-dependent CRISPR-Cas gene editing relies on RNA-guided DNA cleavage that drives cellular machinery to incorporate intended edits. Beyond canonical cleavage, CRISPR nucleases can also execute non-canonical cleavage on the displaced DNA strand. Here, we report a modified design principle for prime editing that leverages the non-canonical cleavage activity, enabling editing in the 5’ direction of a classic nick site. We elucidate the non-canonical-cleavage-driven reverse transcription pathway that initiates upstream of the classic nick site. We identify competition between the canonical and non-canonical routes, which can be modulated by rationally designing RNA templates carrying the intended edits. We demonstrate that cellular physiology enhances editing efficiency from individual reverse-transcription product yet exerts limited influence on pathway competitions. Our findings revise the design principle of prime editing and establish a new dimension for engineering CRISPR-Cas systems by harnessing its intrinsic, non-host-specific, non-canonical cleavage activity.

## Introduction

Biological systems leverage engineered and natural machineries for designed functionalities, while their full capacities are sometimes concealed due to the limited mechanistic understanding. CRISPR-Cas systems have been revolutionizing biological science, technology, and engineering, and their potential has been continuously expanded with the increasingly in-depth understanding of their biology and chemistry. For instance, deploying exclusively the binding properties of Cas proteins enables manipulation of gene expression^1, 2^, selectively inactivating one of the nuclease domains allows intended single-strand DNA cleavage for single-nucleotide resolution gene editing with combined functional enzymes, such as deaminase^3, 4^ and glycosylase^5^, and harnessing the collateral cleavage feature makes DNA or RNA detection possible^6, 7^.

Prime editing is a disruptive gene editing method, providing new horizons for next-generation medicine and agriculture^8–10^. It was invented by harnessing a Cas9 nickase (nCas9) with functional RuvC fused to reverse transcriptase (RT), with a prime editing guide RNA (pegRNA) that integrates a primer binding site (PBS) and a reverse transcription template (RTT) into the single guide RNA scaffold^11^. This design principle allows the intended edits reverse transcribed from the RNA template to be incorporated into targeted loci with high precision and versatility. Many efforts have been made to improve the system for higher efficiency and an expanded editing spectrum by engineering effectors^12–15^, tailoring pegRNAs^16–18^, and orchestrating host determinants, such as dNTP levels^13^, exonuclease^15^, and the mismatch repair (MMR) machinery^19^. These technological advancements have significantly benefited from the biology of CRISPR-Cas systems, and vice versa.

Despite these advances, all reported prime editing systems employ the nicked DNA strand as the primer and thereby restrict editing to the 3’ direction from the nick site, three nucleotides away from the protospacer adjacent motif (PAM), on the cleaved DNA strand (**Fig. 1a**). To edit the DNA upstream of a specific nick, additional designs are necessary to locate the target downstream of another nick site and thus comply with the established design principles. For instance, EXPERT generates an additional nick upstream of the target on the same strand^18^, and rPE^20^, ciPEs^21^, and ActRNA:t system^22^ individually generate additional nick sites with functional HNH to locate the target position downstream of the exposed 3’-hydroxyl group on the complementary strand. Prime editing in the 5’ direction of the nick is considered theoretically impossible through the well-established reverse transcription pathway (**Fig. 1a**). Beyond its canonical endonucleolytic activity, RuvC can also mediate non-canonical cleavage at variable positions upstream of the classic site on the displaced DNA strand in the R-loop. This feature, with the precise cleavage of HNH, has been demonstrated to generate staggered cleavage and is essential in template-free genetic variations in eukaryotic cells^23–25^. Although not previously been repurposed for gene editing, the non-canonical cleavage activity of RuvC provides an opportunity to revise the established design principles of prime editing by generating a DNA primer upstream of the classic nick site, thereby initiating a new reverse transcription pathway and enabling previously impossible editing (**Fig. 1b**).

**Fig. 1.**
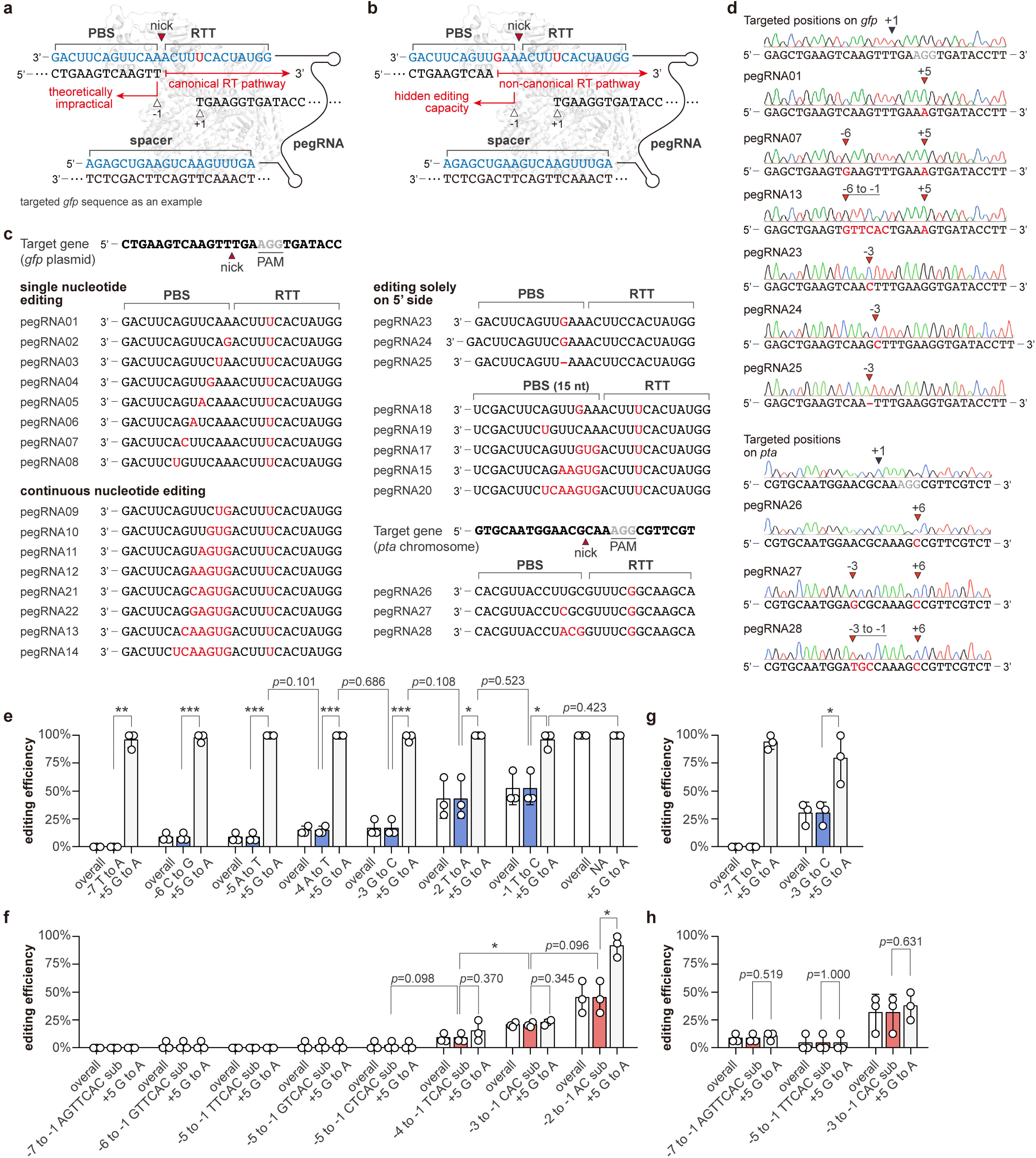
Prime editing in the 5’ direction of a nick. (**a**) Mechanical scheme of canonical prime editing. Reverse transcription extends from position +1 in the 3’ direction of the nicked DNA strand to synthesize a new DNA strand containing intended edits using RTT as the substrate. The first base downstream of the nick is position +1. (**b**) Upstream prime editing through a non-canonical reverse transcription pathway. (**c**) Designs of pegRNAs targeting *gfp* and *pta* with single-nucleotide substitution or continuous substitutions in the 5’ direction of the nick. (**d**) Representative sequencing results of the upstream prime editing in *gfp* and *pta*. Successful edits are highlighted at the designed positions. (**e**) Prime editing efficiencies for single-nucleotide substitution and (**f**) continuous substitutions with a 13-nt PBS. (**g, h**) Prime editing efficiency with a 15-nt PBS. Each pair of blue (or red) and light grey columns indicates upstream and downstream editing, respectively. The overall editing efficiencies (white bars) are consistent with the efficiencies of upstream editing, as all upstream edits are only identified together with the downstream edits. Blue color indicates single-nucleotide substitutions in **e** and **g**, and Red color indicates continuous substitutions in **f** to **h**. The edited positions were indicated by the arrows, and edited bases were highlighted in red. Data are presented as mean values ± SD (n = 3 biological replicates). Statistical significance is determined by *t*-test (*, *p*<0.05; **, *p*<0.01; ***, *p*<0.001, unpaired and two-sided), and the exact *p*-values are provided in Source Data. Source data are provided as a Source Data file. Abbreviations: PBS, primer binding site; RTT, reverse transcription template; pegRNA, prime editing guide RNA.

Here, we report a modified design principle for prime editing that leverages the non-canonical RuvC cleavage activity, allowing a basic prime editing system to modify positions in the 5’ direction of the classic nick site. We systematically characterize the hidden capacity of prime editing and elucidate the non-canonical-cleavage-driven reverse transcription pathway that initiates upstream of the classic nick site. We demonstrate the competition and modulation between the canonical and non-canonical routes, and identify strategies to enhance editing efficiency. Our findings revise the current understanding and design principle of prime editing by integrating of the non-canonical cleavage activity of Cas9 into precision gene editing.

## Results

### Enabling prime editing in the 5’ direction of a classic nick site

To uncover the untapped capability of prime editing through the non-canonical cleavage, we customized PE2^11^ for *Escherichia coli* with only essential components, including the fusion of *Streptococcus pyogenes* nCas9 and engineered Moloney murine leukemia virus (M-MLV) RT as effector, and the pegRNA containing designed edits (**Supplementary Fig. 1**). We targeted a plasmid-carried *gfp* to avoid misleading conclusions due to the low efficiency of prime editing (**Fig. 1c**). First, we designed pegRNAs with a single-nucleotide substitution in the PBS with an edit in the RTT serving as a positive control indicating the efficacy of prime editing (**Fig. 1c**). With the first position downstream of the nick as position +1, the editing positions ranged from –1 to –7 (**Fig. 1c**), leaving at least 6 nt of a 13-nt PBS as the binding site^11^. While part of the PBS functionally served as the RTT in these designs, we retained the terminology to illustrate the results and mechanism. Successful editing was observed from positions –1 to –6, while position –7 could not be modified (**Figs. 1d, 1e and Supplementary Fig. 2a**). Next, we designed pegRNAs containing edits with continuous substitutions ranging from 2 to 7 nt (**Fig. 1c**), and a maximal 6-nt substitution (–6 to –1 GTTCAC) was achieved (**Figs. 1d, 1f and Supplementary Fig. 2b**). Due to potential sequence preferences^11^, the success of 5-nt editing was observed with varied editing sequences at position –5 (–5 to –1 GTCAC and CTCAC) instead of the original design (–5 to –1 TTCAC) (**Figs. 1c, 1f and Supplementary Fig. 2b**). Besides substitutions, we also achieved continuous 5– and 6-nt insertions and deletions upstream of the classic nick, respectively (**Supplementary Fig. 3**). These results further demonstrate that the exposed 3’-hydroxyl group functions similarly to that generated by the canonical cleavage and leads to reverse transcription to incorporate all types of edits from the RNA template. Taking position –3 as an example, we found that the editing could be achieved alone without accompanying edits in RTT, and all three types of editing, substitution, insertion and deletion, were possible (**Figs. 1c and 1d**). In addition, we targeted the *pta* gene in the chromosome of *E. coli* (**Fig. 1c**), and, as designed, successful editing was identified (**Fig. 1d**).

These results demonstrated that prime editing can reach out to position –6 with a 13-nt PBS. Previous studies reported that a minimal length of binding site is required to anchor the RNA template to the target DNA strand, and shorter lengths reduce editing efficiency^11, 26^. The maximal capacity we observed might be underestimated due to the significantly shorter PBS available for binding. As such, we elongated the PBS to 15 nt to mitigate the impacts of the binding site (**Supplementary Fig. 4a).** Though single-nucleotide substitution at position –7 still failed (**Fig. 1g and Supplementary Fig. 4b**), the –7 to –1 AGTTCAC substitutions and the previously unsuccessful –5 to –1 TTCAC substitutions were obtained (**Fig. 1h and Supplementary Fig. 4b**). This maximal capacity is in agreement with previous reports that the cleavage site can reach position –7 on the displaced DNA strand^24, 27^. As a common observation of mixed sequencing signals after base editing and prime editing in bacteria, pure edited strains can be obtained through an additional round of segregation^28, 29^. We characterized the fidelity of the edited strains after an additional round of segregation, and identified clean editing in all resulting strains as shown in corresponding sequencing chromatograms. In addition, we designed two more pegRNAs targeting different positions on the gene and similar observations were identified (**Supplementary Fig. 5**).

### The non-canonical pathway competes with the canonical prime editing route

Following the established design principles, nCas9 cleaves on the displaced single-strand DNA within the formed R-loop, generating a RuvC nick three nucleotides from the PAM (**Fig. 2a**). The cleaved DNA strand binds to the PBS and serves as a DNA primer. The canonical reverse transcription pathway (cRTP) then initiates at this classic nick site and incorporates the designed edits from RTT (**Fig. 2a**). As such, cRTP in theory cannot mediate editing in the 5’ direction of the nick site. When an edit is designed in the PBS, the non-canonical cleavage activity may generate an exposed 3’-hydroxyl group at or farther than the mismatched locus between the DNA and RNA (**Fig. 2a**), thereby priming a non-canonical reverse transcription pathway (nRTP) and implementing the designed edits from PBS (**Fig. 2b**). This mechanism was further demonstrated *in vitro* using purified prime editing effectors in defined reaction systems. By supplying distinct pegRNAs, we observed reverse-transcription products containing edits from RTT, PBS or both (**Supplementary Fig. 6**), which were in agreement with our *in vivo* observations. The results also exclude the requirements for host-dependent factors that may interact with the prime editing system and, theoretically, making the revised design principle generalizable for diverse hosts.

**Fig. 2.**
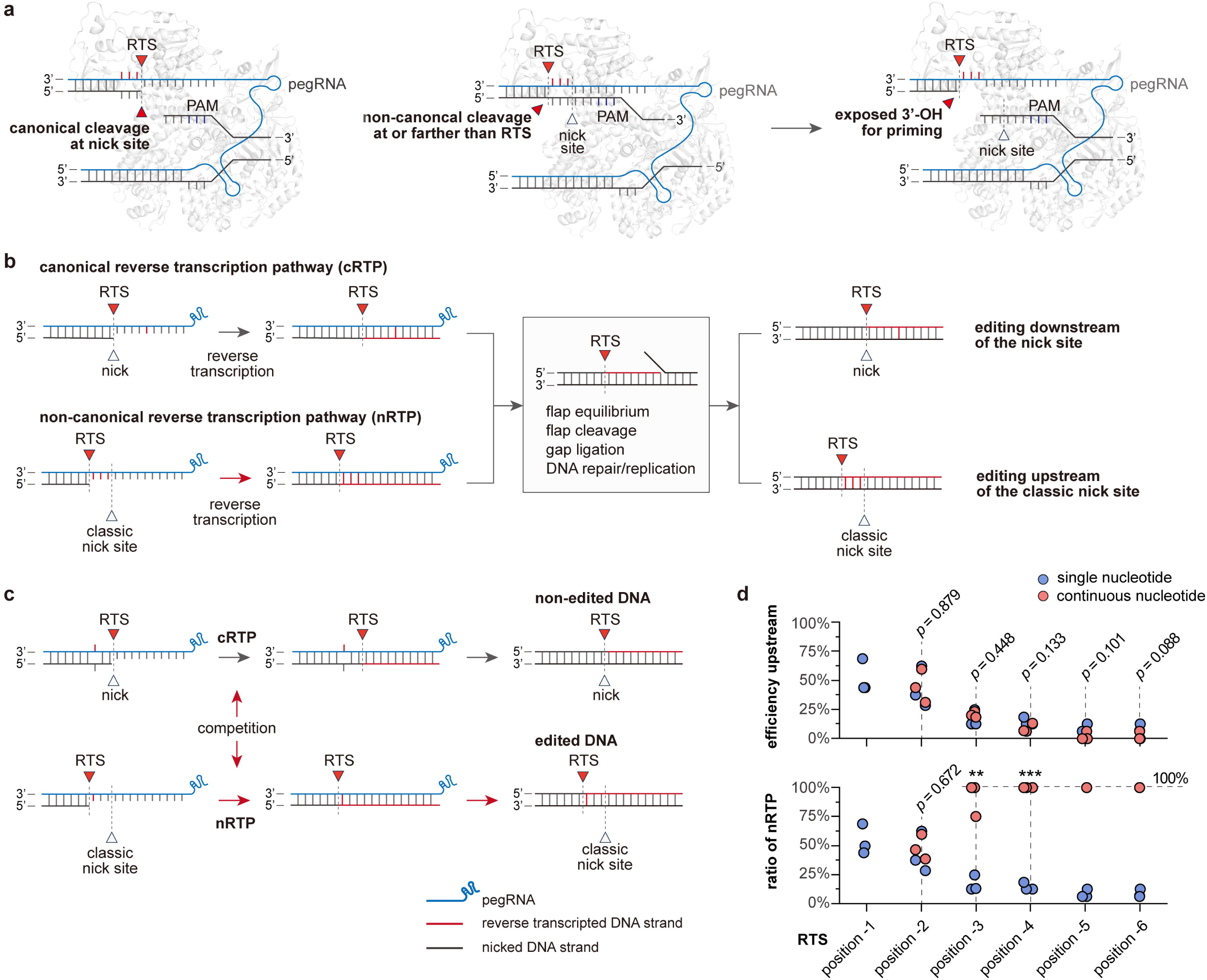
Non-canonical-cleavage-driven reverse transcription and pathway competitions. (**a**) The canonical and non-canonical cleavage of Cas9 that generates exposed 3’-hydroxyl groups at distinct loci for priming. (**b**) Mechanistic illustration of cRTP and nRTP. In the cRTP, reverse transcription initiates at the classic nick site, 3 nucleotides from the PAM, while, in the nRTP, the non-canonical cleavage creates a distinct RTS with an exposed 3’-hydroxyl group upstream of the classic nick site, priming the non-canonical reverse transcription. Both pathways produce reverse-transcribed DNA containing the designed edits, which are integrated into the target position through subsequent DNA processing. (**c**) Competition between the cRTP and nRTP. The editing outcome is determined by the pathway and only nRTP leads to editing upstream of the classic nick site. (**d**) Quantitative analysis and modulation of pathway competition. The ratio of nRTP to cRTP is shown for single-nucleotide (red-filled circle) and continuous-nucleotide (red-filled circle) substitutions at specific RTSs from –1 to –6, and longer continuous nucleotide substitutions enhance the competency of nRTP but reduce the overall editing efficiency. Data are presented as mean values ± SD (n = 3 biological replicates). *p*-values indicate statistical significance between the two pathways at each position and are determined by t-test (*, *p*<0.05; **, *p*<0.01; ***, *p*<0.001, unpaired and two-sided). The exact *p* values are provided in Source Data. Source data are provided as a Source Data file. Abbreviations: pegRNA, prime editing guide RNA; RTS, reverse transcription start site; cRTP, canonical reverse transcription pathway; nRTP, non-canonical reverse transcription pathway; PAM, protospacer adjacent motif.

As the classic nick site is no longer necessarily the starting position of reverse transcription, we introduce the term reverse transcription start site (RTS), which can be generated by either canonical or non-canonical cleavage, to interpret the revised design principle (**Fig. 2a**). We observed that editing upstream of the nick always accompanied edits on the 3’ side when edits on both sides were designed, as supported by the overall editing efficiencies and the separately reported efficiency for each side (**Figs. 1d – 1h, and Supplementary Table 1**). No standalone editing upstream of a nick was identified, suggesting that a single reverse-transcribed DNA strand from the RTS orchestrates an individual editing event. Interestingly, for single-nucleotide substitutions upstream of the nick, the editing efficiencies downstream of the nick were mostly around 100%, regardless of those upstream of the nick, from 87.50% to 100.00% (**Fig. 1e**). These results suggested both pathways were functional yet competing with each other, leading to distinct editing outcomes, and only if the editing goes through nRTP, the intended edits upstream of the nick can be integrated (**Fig. 2c**). In these cases, while the non-canonical cleavage can initiate nRTP, cRTP is still likely to occur at the classic nick site, as a single mismatch may not completely disrupt the RNA-DNA duplex. We calculated the thermodynamics of duplex formation between the PBSs of different pegRNAs and their target DNAs using nearest neighbor parameters^30, 31^. The results showed that the actual PBS-DNA duplexes in cRTP and nRTP are comparably thermodynamically favorable, while cRTP becomes more favorable as the intended edit moves farther from the classic nick site (**Supplementary Table 2**). Differently, the editing efficiencies of the 3– to 7-nt continuous substitutions upstream of the nick were similar to those downstream of the nick (**Figs. 1f and 1h**), implying that nRTP was the predominant pathway, whereas cRTP was rare. This can also be partially explained by the thermodynamics, as nRTP becomes more favorable with longer continuous substitutions (**Supplementary Table 2**). Notably, the thermodynamics may also contribute to the unique editing profile of the –1 to –2 substitutions (**Fig. 1f and Supplementary Table 2**).

Meanwhile, we found that the editing efficiencies of continuous substitutions decreased with increasing distance between RTS and the nick (**Fig. 1f**), and a similar decrease in editing efficiencies upstream of the classic nick was identified for single-nucleotide substitutions (**Fig. 1e**). This can be interpreted by a combined effects of shorter actual binding sites and the non-canonical cleavage activity distant from the nick. Quantitatively, we found that editing efficiencies with a specific RTS were similar, implying a relatively determined competency of the non-canonical cleavage at a certain RTS (**Fig. 2d**). Therefore, the competition between cRTP and nRTP reflects the combined effects of the ability of RuvC to mediate the canonical or non-canonical cleavage and the binding properties, including the thermodynamics, of the formed RNA-DNA duplexes that actually initiate reverse transcription at different positions. As the binding properties can be modulated by the PBS sequences, we underscore that editing outcomes can be orchestrated by rationally designing continuous nucleotide modifications to minimize cRTP from the classic nick site.

### Disrupting 3’-exonuclease and mismatch repair system enhances editing from single reverse transcripts

Host physiology is essential for prime editing, especially after generating the reverse-transcribed DNA containing the edits. Exonucleases have two distinct impacts that 3’ exonuclease cleaves the 3’ flap leading to decreased editing efficiency^32^, while 5’ exonuclease cleaves the 5’ flap, increasing editing efficiency^15^ (**Fig. 3a**). As such, we employed deamination-based CRISPR-Cas base editing and inactivated *sbcB* alone, the gene encoding the endogenous 3’ exonuclease DNase EXO I, as well as *sbcB*, *xseA*, and *exoX* together, generating two engineered hosts with impaired 3’ exonuclease activity (**Supplementary Fig. 7**). Taking plasmid-carried *gfp* as a target, we identified significantly increased efficiency of single-nucleotide substitution at position –3 in the *sbcB*-deficient strain, while a marginal rise in the strains with three inactivated genes (**Fig. 3b**). Notably, the editing efficiencies downstream of the nick were still approaching 100% (**Fig. 1e and 3b**). This may be due to the preference of *sbcB* for flapped DNAs of different lengths^33, 34^, leading to the differential modulation of DNA flaps through the two distinct pathways. As expected, the efficiency of continuous substitutions was significantly enhanced, reaching 91.67% in the *sbcB*-inactivated strain and 97.78% in the triple-inactivated strain (**Fig. 3b**). These results indicate that the inactivation of 3’ exonuclease can modulate the equilibrium of flap cleavage and retain more reverse-transcribed DNA from nRTP, and slightly regulate the editing outcomes after pathway competitions.

**Fig. 3.**
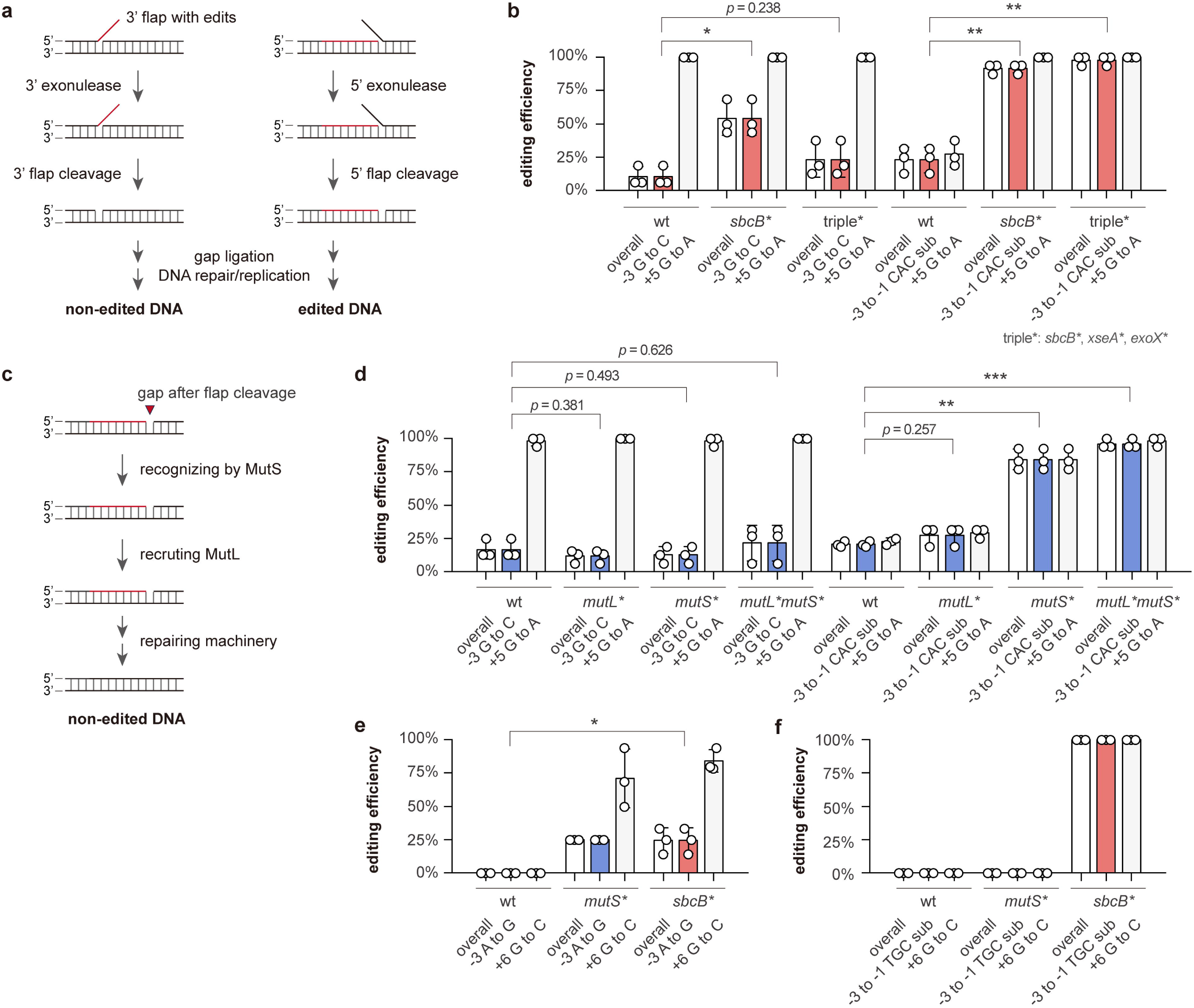
Host physiology enhances prime editing efficiency upstream of the nick. (**a**) Schematic illustration of flap cleavage processing by exonucleases and (**c**) mismatch repair mechanism (MMR). After reverse transcription, the 3’ flap containing the edited sequence can be cleaved by 3’ exonuclease (**a**), leading to loss of the intended edits. Once the flap equilibrates, the remaining gap can be recognized by MutS and mediated by MutL (**c**), leading to the removal of edits through MMR. (**b**) Prime editing efficiencies of *gfp* in the wild-type (wt) strains with 3’-exonuclease-deficient strains, and (**d**) with MMR-deficient strains. The plasmid-carried *gfp* was selected as a target and RTS at –3 as a representative. (**e**) Editing efficiencies in chromosomal *pta* for single-nucleotide substitution and (**f**) continuous substitutions in wild-type, MMR-deficient (*mutS\**) and 3’ exonuclease-inactivated (*sbcB\**) strains. Each pair of colored (blue or red) and light grey columns indicates upstream and downstream editing, respectively. The overall editing efficiencies (white bars) are consistent with the efficiencies of upstream editing, as all upstream edits are only identified together with the downstream edits. Blue color indicates MMR-deficient strains in **d** and **e**, and Red color indicates 3’-exonuclease-deficient strains in **b, e** and **f**. Data are presented as mean values ± SD (n = 3 biological replicates). Statistical significance is determined by t-test (*, *p*<0.05; **, *p*<0.01; ***, *p*<0.001, unpaired and two-sided), and the exact *p* values are provided in Source Data. Source data are provided as a Source Data file.

After flap cleavage, the left gap can be recognized by mismatch repair (MMR) systems (**Fig. 3c**), and, debatably, manipulating MMR may lead to higher editing efficiencies^19, 35^. Leveraging base editing, we deactivated the key genes encoding the MMR individually by installing early STOP codons (**Supplementary Fig. 8**). Disruption of MMR exerted limited effects on single-nucleotide substitutions upstream of the nick, maintaining an efficiency around 15.18%, while the edits downstream of the nick approached 99.31% (**Fig. 3d**). These editing profiles agreed with our discovery that editing efficiencies at a specific RTS are similar (**Fig. 1e**), illustrating that the competition between nRTP and cRTP could hardly be modulated through MMR. To the contrary, the 3-nt continuous substitutions were significantly enhanced in *mutS-* and *mutLS-* deficient strains, reaching a maximum of 95.83%, suggesting that 75.22% of the edits from single reverse transcripts were removed by MMR in the wild-type strain (20.61%) (**Fig. 3d**).

Next, we evaluated the effects of these two mechanisms on chromosomal editing. We targeted the *pta* gene in *sbcB-* and *mutS-*deactivated strains, respectively. The two individual gene inactivations showed similar enhancements compared to multi-gene inactivations (**Fig. 3e and 3f**). We observed that single-nucleotide substitution efficiency could be significantly increased in these two engineered hosts, while no desired edits were identified in the wild-type strain under our current experimental setting. The editing at position –3 was around 24.36%, while that at position +6 approached 76.98% (**Fig. 3e**). For continuous substitutions, significantly increased editing efficiency could only be observed in the *sbcB*-inactivated strain, with no detectable edits in the wild-type or *mutS* mutant (**Fig. 3f**). As only reverse-transcript though nRTP existed in these 3-nt editing events, 3’ exonuclease might be a more determinant factor than MMR in orchestrating the non-canonical prime editing via the nRTP. Taken together, exisiting strategies, such as modulating 3’-exonuclease activity and MMR, can enhance prime editing driven by an individual reverse transcription pathway, and the revised design principle is unlikely to affect these strategies, as they mainly function after reverse transcription has generated the DNA containing the intended edits. Yet, these strategies are difficult to modulate pathway competitions during reverse transcription.

### Non-canonical prime editing creates previously impossible edits

Our findings are not just a conceptual disruption but also allow previously impossible modifications in the genome. We evaluated the phenotype of the strains containing the edited *gfp*. As expected, insertion and deletion at position –3 led to inactivation of *gfp* through open reading frame (ORF) shifts (**Supplementary Fig. 9**). Interestingly, we found the substitution of G-to-C at the same locus showed only lower fluorescence due to a lysine to asparagine missense mutation (Lys113Asn), which was similar to that resulting from a +5 G-to-A editing leading to the Gly116Ser mutation (**Supplementary Figs. 9 and 10**). This G-to-C change cannot be easily achieved following the conventional design principle of prime editing, nor deamination-mediated base editing due to the editing capability and PAM requirement. To further demonstrate the potential, we designed edits in two more genes in the genome, leading to the gain and loss of functions that could not be achieved before. First, we designed prime editing to install an early STOP codon into *glnA*, encoding the glutamine synthetase, which catalyzes the synthesis of glutamine (**Fig. 4a**). By doing so, we aimed to generate an auxotrophic strain that cannot grow in minimal medium without glutamine. We chose to change the codon ACT for Thr25 directly to the STOP codon TAG (**Fig. 4b**), and successful editing and the expected phenotypic changes were observed (**Figs. 4c and 4d**), demonstrating the functionality of our new design principle. Although early STOP codons can be installed at different loci with other methods, this specific editing could not be obtained without the revised prime editing principle, as the target is located both upstream and downstream of the nick (with the nearest AGG PAM, **Fig. 4b**). This has not yet been documented in previous reports.

**Fig. 4.**
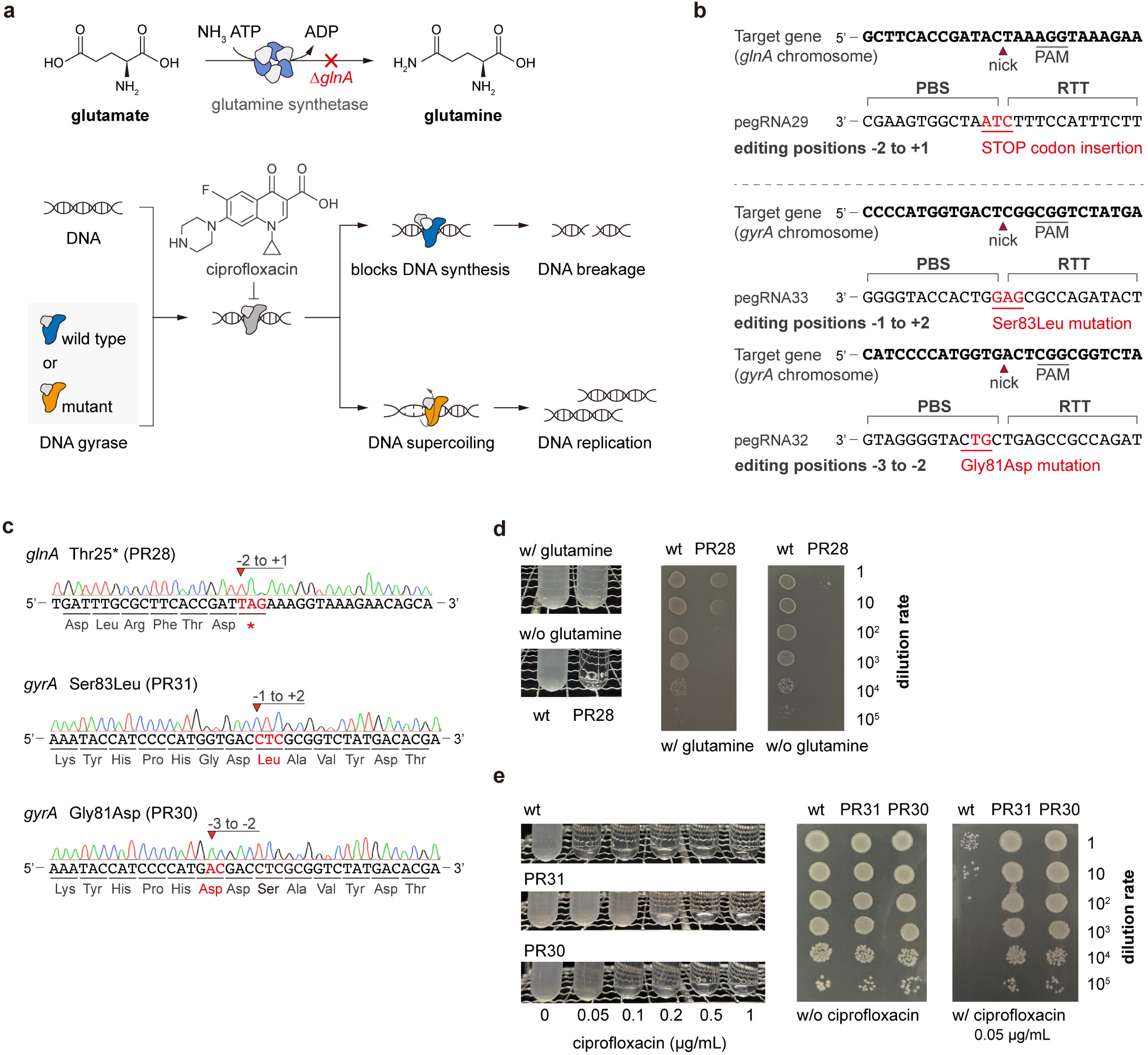
Non-canonical prime editing enables modifications beyond conventional design limits. (**a**) The *glnA*-mediated glutamine metabolism and *gyrA*-mediated ciprofloxacin tolerance. Disruption of *glnA* renders an auxotrophic strain that cannot grow without glutamine, and mutations of *gyrA* at amino acid positions 83 and 81 alter the DNA gyrase active site, thereby reducing the binding affinity with ciprofloxacin. (**b**) Designs of pegRNAs for *glnA* and *gyrA* editing in the 5’ direction of the nick, leading to loss and gain of functions, respectively. (**c**) Sequencing results of *glnA* and *gyrA* with intended edits upstream of the nick. (**d**) Phenotypical evaluation of the *glnA* inactivated strains and (**e**) ciprofloxacin susceptibility test of *gyrA* mutant strains with different levels of ciprofloxacin in liquid medium and spotted onto agar plates with 0.05 μg/mL ciprofloxacin. The edited positions were indicated by arrows, and the altered amino acids and edited bases were marked in red.

As a second example, we designed edits in the *gyrA* gene, which encodes one of the essential subunits of DNA gyrase (**Fig. 4a**). As reported, a serine-to-leucine substitution at position 83 (Ser83Leu) or a glycine-to-aspartic acid mutation at position 81 (Gly81Asp) will grant the mutated strain higher tolerance to ciprofloxacin^36^. We admit that these two modifications can be made with a conventional design by elaborately choosing PAMs. For instance, the Ser83Leu mutation can be achieved with the CGG PAM starting at nucleotide locus 248 of *gryA*, and the intended edits would be located from positions +3 to +5 downstream of the target DNA. To demonstrate our new principle, we chose the CGG PAM starting at nucleotide locus 251 instead (**Fig. 4b**), thereby locating the desired editing from –1 to +2 on both sides of the nick. We successfully obtained the desired editing and confirmed the altered phenotype, generating a ciprofloxacin-tolerant strain up to 0.2 μg/mL (**Figs. 4c, 4e and Supplementary Fig. 11**). To obtain the Gly81Asp mutation, we selected the CGG PAM at nucleotide locus 248 of *gryA*, and the designed edits located at position –3 and –2 (**Fig. 4b**). This was a new editing pattern even in the present study, while we did not edit the nucleotide adjacent to the nick in the 5’ direction. The editing was successful and generated another ciprofloxacin-tolerant strain. This strain was more sensitive than the strain with Ser83Leu (**Figs. 4c, 4e and Supplementary Fig. 11**), which was in agreement with previous reports on *gyrA* mutations^36^.

## Discussion

All cleavage-dependent CRISPR-Cas gene editing relies on the single– or double-strand breaks on the targeted DNA, then driving cellular machinery to incorporate intended edits^37^. The non-canonical staggered cleavage of Cas9 has been known and interrogated to interpret and predict CRISPR-mediated template-free mutations^23–25^. Particularly, staggered ends can be generated when the non-canonical activity of RuvC cleaves the displaced DNA strand at variable positions upstream of the classic position, while HNH cleaves the complementary DNA strand at its classic position. Here, we report that the non-canonical cleavage activity of Cas9 expands the design principles of prime editing, enabling previously impossible editing in the 5’ direction of a classic RuvC nick site, three nucleotides away from the PAM. When no edits are designed upstream of the nick, the classic nick site serves as the position where reverse transcription starts and coincides with the RTS, a new term we introduced to illustrate the revised principle. When an edit is designed upstream of the nick, the non-canonical cleavage activity of RuvC may generate RTS other than the classic nick site, where DNA sequence mismatches the RNA template in the PBS, leading to a non-canonical reverse transcription pathway (**Figs. 2a and 2b**). While we base our revised design principle on the non-canonical RuvC activity, it remains debated whether the exposed 3’-hydroxyl group farther than position –3 is generated by direct non-canonical cleavage^24^ or by subsequent trimming following an initial cleavage^27^. Although our observed cleavage positions are consistent with both mechanisms, we cannot distinguish the origin of the 3’-hydroxyl group since reverse transcription initiates only after its formation. Notably, this distinction does not affect our revised principle of prime editing, as either mechanism generates an exposed 3’-hydroxyl group that can initiate reverse transcription upstream of the classic nick site.

As a reverse transcription pathway, nRTP is similar to cRTP and can introduces substitutions, insertions, and deletionse. By interrogating the roles of 3’-exonuclease activity and MMR, we infer that nRTP primarily affects the reverse-transcribed DNA, making it not only applicable to other prime editing systems but also compatible with existing strategies to enhance editing efficiencies. However, nRTP is not dominant yet competes with cRTP. This competition reflects the integrative effects of the intrinsic capabilities of the prime editing system, PE2 in this study, in generating canonical and non-canonical cleavage and in forming stable RNA-DNA duplexes to prime reverse transcription, thereby constraining the editing outcomes once the edits are decided. A promising approach to enhance the competency of nRTP is to rationally design the intended modifications in PBS, and a longer length of modified nucleotides would reduce the probability of cRTP. In return, by elaborately designing RNA and DNA mismatch patterns, we may be able to tune the position of the non-canonical cleavage.

Gene editing approaches are critical for biological science, technology and engineering. Prime editing, in particular, can perform all 12 types of nucleotide substitutions and small indels in diverse living organisms^9^. It has been employed to interrogate and treat human diseases^38, 39^ and to engineer smarter plants for more sustainable human societies^40–42^. Besides, it has been adapted for microbes to make precision gene editing at single-nucleotide resolutions, such as in probiotics^28^ and pathogens^43, 44^. The necessity of such edits is boosting due to the massive available omics data and information generated from pooled libraries^45^ and adaptive laboratory evolutions^46^. Recreating desired biological features requires only countable modifications with high precision. For instance, three mutations in *pgi*, *crp* and *rpoB* are already enough to engineer a superior artificial autotroph^47^, and two mutations in *rpoC* plus one in the transcriptional regulator are adequate to generate a faster-growing formatotroph^48^. With its naturally versatile editing capacity, the spectrum that prime editing can achieve is crucial for further expanding its utilizations. By revising the design principle, we show that a basic prime editing system, like PE2, can access positions upstream of a classic nick site, and the desired edits can be introduced by simply implementing a change in PBS rather than RTT. However, the prime editing outcomes, including the efficiency and maximal length of edits, reflects the combined effects of reverse transcription and subsequent cellular processing, thereby requiring further endeavors to establish such systems in a specific living organism. As non-canonical cleavage is an intrinsic feature of CRISPR nucleases rather than host-specific cellular physiology, we envision more gene editing innovations that leverage this unusual capacity in living organisms with functional CRISPR-Cas systems.

## Methods

### Strains and media

*E. coli* DH5α (Takara) was used for molecular cloning in general. *E. coli* MG1655 was employed as the wild-type strain, and it was transformed with a *gfp*-carrying plasmid, generating PR19, for evaluating the unknown competency of prime editing. To evaluate the role of DNA exonuclease and DNA mismatch repair (MMR), we constructed DNA exounuclease-deficient strains PR20 (*sbcB* Gln129*) and PR21 (*sbcB* Gln129*, *xseA* Gln116*, and *exoX* Trp63*), and MMR-deficient strains PR22 (*mutL* Gln4*), PR23 (*mutS* Gln138*), and PR24 (*mutL* Gln4* and *mutS* Gln138*). These mutated strains were then transformed with the *gfp*-carrying plasmid as evaluation platforms for either plasmid or chromosome editing. All strains used and generated in this study are listed in **Supplementary Table 3.** *E. coli* strains were grown in Luria-Bertani (LB) medium (10 g/L tryptone, 5 g/L yeast extract, 10 g/L NaCl) and selected on LB solid medium (with 1.5% agar). Carbenicillin, as a stable alternative for ampicillin (100 µg/mL) and gentamicin (20 µg/mL) were supplemented for plasmid selection and maintenance. Isopropyl-β-D-thiogalactoside (IPTG) was added to a final concentration of 1 mM to induce prime editing and base editing. Chemically competent cells were prepared using a classic CaCl_2_ protocol. For phenotypic analysis, wild-type and edited strains were grown in M9 minimal medium containing 15.14 g/L Na_2_HPO_4_·12H_2_O, 3.0 g/L KH_2_PO_4_, 0.5 g/L NaCl, 1.0 g/L NH_4_Cl, 0.241 g/L MgSO_4_, 0.011 g/L CaCl_2_ and 4 g/L glucose. Appropriate amino acids (5 mM L-glutamine) or various concentrations of ciprofloxacin (0.05, 0.1, 0.2, 0.5, and 1 μg/mL) were supplemented when required.

### Plasmid construction

General DNA manipulation and molecular cloning were conducted as follows. PCR amplification of DNA fragments was performed using PrimeSTAR^®^ Max DNA Polymerase (Takara), and plasmid assembly was achieved via the In-Fusion^®^ Snap Assembly Master Mix (Takara). Plasmids were extracted using the QIAprep Spin Miniprep Kit (QIAGEN) and DNA purification was carried out using the QIAquick PCR Purification Kit (QIAGEN). Gel electrophoresis and Sanger sequencing were employed for plasmid validation. DNA concentration was quantified by NanoDrop One (Thermo Scientific).

To investigate the capacity of prime editing, we constructed the target plasmid pGFP using pBR322 as a backbone with the pBBR1MCS-5 oriV and pBBR1MCS-5 rep and *bla* as the selection marker. The *gfp* gene from pAM4787^49^ (Addgene #120088, generously provided by Dr. Susan Golden’s laboratory) was incorporated as the gene of interest. Next, we customized PE2 for *E. coli*, employing a previously established prime editing system, pPE.S, containing the engineered Cas9 nickase (nCas9, H840A) from *Streptococcus pyogenes* fused to an *E. coli* codon-optimized M-MLV reverse transcriptase (RT) under the control of a *lacI*-P_trc_ inducible system^28^. The *oriR101* in pPE.S was replaced with the *p15A* ori for better performance, generating the working plasmid template pRC. pRC-His was subsequently constructed based on pRC by adding a 6 × His tag into the PE2 expression cassette for protein purification. All pegRNAs were commercially synthesized on a pUC57 plasmid driven by a constitutive promoter P_J23119_. Then, the pegRNA cassettes were amplified and incorporated into pRC to generate the pRC12 – pRC47 serial editing plasmids. To investigate the role of DNA exonucleases and MMR in prime editing, we employed the established *E. coli* cytosine base editing system, pBeCas9^29^, to inactivate key genes encoding DNA exonucleases and MMR, respectively. The system contains dCas9, PmCDA1, *ugi* and an LVA tag under the control of a *lacI*-P_trc_ inducible system, along with a temperature-sensitive origin of replication *oriR101* and *bla* as the selection marker. The P_J23119_-driven gRNA cassettes were generated via inverse PCR and were incorporated into pBeCas9 via gene assembly, generating base editing working plasmids. All pegRNA sequences provided in **Supplementary Table 4,** gRNA sequences in **Supplementary Table 5**, and plasmids are provided in **Supplementary Table 6**. All primers used in this study are listed in **Supplementary Table 7.**

### Mutating DNA exonucleases and MMR with base editing

We employed deamination-mediated base editing to disrupt DNA exonuclease genes, including *sbcB*, *xseA* and *exoX* (**Supplementary Table 8**), by introducing early stop codons. Specifically, the Gln129 in *sbcB* (encoded by CAG), Gln116 in *xseA* (encoded by CAG) and Trp63 in *exoX* (encoded by TGG) were converted to STOP codons (TAG or TAA) with customized gRNAs (gRNA-sbcB, gRNA-xseA and gRNA-exoX, **Supplementary Table 5**). To construct the mutant strains, wild-type MG1655 was first edited with pBeCas9-sbcB to generate PR20. The strain was subsequently edited with pBeCas9-xseA and then with pBeCas9-exoX to yield the final triple-inactivated strain PR21 (**Supplementary Tables 3 and 6**). Base editing was also employed to disrupt MMR by introducing early STOP codons into *mutS* and *mutL* (**Supplementary Table 8**). The Gln4 in *mutL* and Gln138 in *mutS* (both encoded by CAG) were converted to STOP codons (TAG) with specifically designed gRNA-mutL and gRNA-mutS (**Supplementary Table 5**). To generate the mutated strains, the wild-type MG1655 was edited with pBeCas9-mutL and pBeCas9-mutS, respectively, to generate PR22 and PR23. PR24 was constructed by editing PR22 with pBeCas9-mutS (**Supplementary Tables 3 and 6).** All base editing was transformed with 25 ng of plasmid DNA via heat shock. Then, the cells were recovered in LB at 30 °C for 1 h, followed by induction with IPTG for 4 h at 30 °C. The edited strains were selected on solid media with appropriate antibiotics. Successful editing was confirmed by Sanger sequencing.

### Design of pegRNAs

We designed pegRNAs targeting *gfp* on pGFP and *pta* in the chromosome to evaluate the capacity of prime editing. The pegRNAs contained a 13-nt PBS and 13-nt RTT unless otherwise specified. Targeting the plasmid-carrying *gfp*, single-nucleotide substitutions from positions –1 to –7 (pegRNA02 – pegRNA08) and continuous substitutions ranging from 2 to 7 nt (pegRNA09 – pegRNA14, pegRNA21 and pegRNA22) were designed with an editing at position +5 in RTT to indicate the efficacy of prime editing. pegRNA01 was designed for +5 substitution. Since the initial design for 5-nt continuous substitutions (pegRNA12, –5 to –1 TTCAC) failed, we designed two alternative pegRNAs (pegRNA21, –5 to –1 GTCAC; pegRNA22, –5 to –1 CTCAC) for 5-nt continuous substitutions. Next, we designed pegRNA23, pegRNA24 and pegRNA25 for gene substitution, insertion, and deletion at position –3 without including edits in RTT. PegRNA 34 and pegRNA35 were designed for 3-nt continuous substitutions at two additional *gfp* sites. PegRNA36 – pegRNA39 were designed for 5-nt insertion, 6-nt insertion, 5-nt deletion and 6-nt deletion, respectively. For *in vitro* prime editing, pegRNA40 was designed for +5 substitution, while pegRNA41 and pegRNA42 were designed for upstream 3-nt continuous substitutions with or without +5 substitution. PegRNA40 – 42 contained the T7 promoter.

Subsequently, we designed pegRNAs with a 15-nt PBS, including pegRNA18 and pegRNA19 for single substitutions at positions –3 and –7, and pegRNA17, pegRNA15, and pegRNA20 for continuous substitutions of 3 nt, 5 nt and 7 nt upstream of the nick. Targeting the chromosomal *pta*, we designed pegRNAs to make +6 substitution (pegRNA26), –3 substitution (pegRNA27) and –3 to –1 continuous substitutions (pegRNA28). PegRNA27 and pegRNA28 also contain edits at position +6. Targeting the chromosomal *glnA*, we designed pegRNA29 to yield –2 to +1 continuous substitutions, and targeting *gyrA*, we designed pegRNAs to make –3 to –2 continuous substitutions (pegRNA32) and –1 to +2 continuous substitutions (pegRNA33). All pegRNAs were commercially synthesized with their sequences provided in **Supplementary Table 4.**

### Prime editing

The wild-type *E. coli* MG1655 and pGFP-containing strain PR19 (**Supplementary Table 3**) were employed as the target strains. For plasmid editing, we transformed PR19 with prime editing plasmids (pRC12 – pRC35) (**Supplementary Table 6),** generating correspondingly edited strains PR36 – PR53, PR69 and PR70 (**Supplementary Table 3**). For chromosome editing, we directly transformed with prime editing plasmids targeting *pta* (pRC36 – pRC38), *glnA* (pRC39) and *gyrA* (pRC40 and pRC41). For DNA exonuclease and MMR evaluation, pRC37 and pRC38 were transformed into pGFP-containing strains (PR54 – PR58) for plasmid editing or directly into PR20 (*sbcB*\*) and PR23 (*mutS*\*) for chromosome editing.

All prime editing was transformed with 25 ng of individual plasmid DNA via heat shock. Cells were recovered in LB medium for 1 h (with carbenicillin for plasmid editing). The cells were then induced for 24 h in fresh LB medium containing 1 mM IPTG and antibiotics (with carbenicillin and gentamicin for plasmid editing, or gentamicin alone for genome editing). When necessary, we extended the induction time for chromosomal prime editing^35^. Successful editing was confirmed by Sanger sequencing of the target region.

### Sanger sequencing

The targeted DNA sequences of randomly selected colonies were amplified via PCR. The PCR products were Sanger sequenced to check the edits. The raw sequencing results were aligned to their respective reference sequences in SnapGene 8.0.3. Chromosomal loci *pta* (ECK2291), *mutS* (ECK2728), *mutL* (ECK4166), *sbcB* (ECK2005), *xseA* (ECK2505), *exoX* (ECK1845), *glnA* (ECK3863) and *gyrA* (ECK2223) were aligned to the *E. coli* K-12 MG1655 reference genome (the National Center for Biotechnology Information accession number NC_000913), while the *gfp* sequence was aligned to a correlated sequence on plasmid pAM4787. All reference sequences are provided in **Supplementary Table 8**.

### *In vitro* reverse transcription experiment

PE2-His was expressed in *E. coli* MG1655 and induced with 1 mM IPTG to the exponential phase (OD_600_ of 0.6). Cells were harvested at 4 °C, washed with PBS and resuspended in a lysis buffer containing protease inhibitor cocktail (P1030, Beyotime). Cell disruption was performed by lysozyme treatment (1 mg/mL) and ultrasonication on ice. After RNase A (10 µg/mL) and DNase I (5 µg/mL) treatment, cell debris was removed by centrifugation at 10000 × g for 30 min at 4 °C. PE2-His was then purified using the His-tag Protein Purification Kit (P2230S, Beyotime) with NTA-Co resin. Purified PE2 protein was stored at –80 °C.

For pegRNA preparation, three synthesized pegRNA fragments (**Supplementary Table 4**) with T7 promoter on a plasmid were linearized with XhoI for 1 h and purified before PCR amplification. The PCR product was used for *in vitro* transcription with the T7 Quick High Yield RNA Transcription Kit (R7016S, Beyotime) at 37 °C for 4 h. The DNA template was removed by DNase I, and the RNA product was purified using an RNA purification kit (Tiangen). RNA concentrations were determined using a NanoDrop spectrophotometer.

For *in vitro* prime editing, a linear dsDNA substrate was first amplified by PCR. Each reaction contained PE2 (30 – 50 nM), pegRNA (2 – 5 µM,), dsDNA (3 – 5 nM), and PE reaction buffer (20 mM HEPES, 100 mM NaCl, 5% glycerol, 3 mM MgCl2, 0.2 mM EDTA, 5 mM DTT and 250 µM each dNTP). Reactions were incubated at 37 °C for 2 h and terminated by incubation at 95 °C for 10 min. Then the products were amplified by a primer pair designed exclusively for the intended reverse-transcription product (**Supplementary Fig. 6**) and analyzed by Sanger sequencing.

### Free energy calculation of the RNA-DNA duplexes

The thermodynamic stability of RNA–DNA duplexes was estimated using the nearest-neighbor model for RNA–DNA hybridization^31^. The Gibbs free energy (ΔG) of each duplex was calculated by summing the contributions of adjacent base-pair steps, with a fixed duplex initiation term. Calculations were based on reported parameters^30^. For adjacent base-pairs with two mismatches for which specific parameters were unavailable, the contribution was estimated as the sum of the average contributions of the corresponding individual mismatches. The calculation matrix is included in Source Data. Lower (more negative) ΔG values indicate more thermodynamically favorable duplex formation.

### Analysis of editing efficiency

After determining the edits through Sanger sequencing, we observed some of the colonies containing mixed sequence signals. This was in agreement with previous reports on prime editing and base editing, while the editing could be purified through one-step segregation^28^. To calculate the efficiency, both pure and mixed sequencing signals were counted as edited. The efficiency was calculated as the percentage of successfully edited colonies among all sequencing colonies (1).

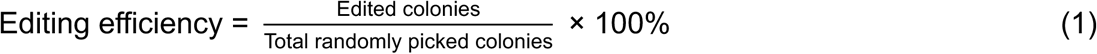

### Plasmid curing

After gene editing, plasmids were cured by culturing the edited strains in antibiotic-free LB medium at 37 °C for 24 h (supplemented with carbenicillin for maintaining pGFP when necessary), followed by streaking on LB agar (or with carbenicillin) plates to obtain individual colonies. Successful plasmid curing was confirmed by determining the plasmid-specific sequences and antibiotic sensitivity to gentamicin.

### Fluorescence measurement

For quantitative fluorescence measurements, cells were harvested by centrifugation, washed twice with phosphate-buffered saline (PBS, pH 7.2), and then diluted to an optical density at 600 nm of 0.01. Fluorescence intensities were subsequently measured using a microplate reader (TECAN). The fluorescence of *gfp* was measured at 488 nm (excitation) and 530 nm (emission).

### Phenotype verification

To verify the functional consequences of the edits, we performed phenotypic verifications of the auxotrophic strain (PR28) and *gyrA* mutants (PR30 and PR31). For the auxotrophic strains, we cultured the wild-type strain and edited strains in M9 minimal medium with or without amino acid (5 mM L-glutamine). For the *gyrA* mutants, we performed a ciprofloxacin susceptibility test with increasing concentrations of ciprofloxacin (0, 0.05, 0.1, 0.2, 0.5 and 1 μg/mL) in liquid and solid media. For assays on plates, all strains were adjusted to OD_600_ = 1, serially diluted from 10^-1^ to 10^-5^, and spotted onto selective plates (M9 with or without glutamine for PR28 and LB with 0.05 μg/mL or 0.2 μg/mL ciprofloxacin for both PR30 and PR31).

## Statistics and Reproducibility

All experiment were conducted with three independent biological replicates (n=3), all of which were included in the analysis. Significance was determined by two-tailed unpaired *t*-test (*, *p*<0.05; **, *p*<0.01; ***, *p*<0.001). The statistical test applied to each comparison are specified in the relevant figure legends. Sample sizes were not determined using a statistical method and no data was excluded. Experimental allocation was not randomized, and the investigators were aware of group assignments during experiments and outcome assessment.

## Data Availability

Source data for editing efficiency, thermodynamic calculation and original images for phenotypical evaluations are provided in the Source Data files. Source data are provided with this paper.

## Author Contributions

P.-F.X. designed the research. P.-R.C. performed the experiments. P.-R.C., X.-Z.Y., and P.-F.X. analyzed the data. P.-R.C. and P.-F.X. drafted the manuscript. X.-Z.Y., S.-G.W., and P.-F.X. revised the manuscript. All authors discussed the results and commented on the manuscript. S.-G.W. and P.-F.X obtained funding for the project.

