## Supplementary information for "Non-canonical cleavage expands prime editing in 5’ direction"

**Supplementary Table 1.** Overall editing efficiencies of intended edits.

| pegRNA | Intended edits [a] | Editing efficiency | | | Host |
|---|---|---|---|---|---|
| | | Repeat 1 | Repeat 2 | Repeat 3 | |
| Targeting at *gfp* | | | | | |
| pegRNA01 | +5 G to A | 100.00% | 100.00% | 100.00% | Wild type |
| pegRNA02 | -1 T to C and +5 G to A | 68.75% | 43.75% | 43.75% | Wild type |
| pegRNA03 | -2 T to A and +5 G to A | 28.57% | 37.50% | 62.50% | Wild type |
| pegRNA04 | -3 G to C and +5 G to A | 25.00% | 12.50% | 12.50% | Wild type |
| pegRNA05 | -4 A to T and +5 G to A | 12.50% | 12.50% | 18.75% | Wild type |
| pegRNA06 | -5 A to T and +5 G to A | 6.25% | 12.50% | 6.25% | Wild type |
| pegRNA07 | -6 C to G and +5 G to A | 6.67% | 12.50% | 6.45% | Wild type |
| pegRNA08 | -7 T to A and +5 G to A | 0.00% | 0.00% | 0.00% | Wild type |
| pegRNA09 | -2 to -1 AC sub and +5 G to A | 31.25% | 43.75% | 60.00% | Wild type |
| pegRNA10 | -3 to -1 CAC sub and +5 G to A | 23.08% | 20.00% | 18.75% | Wild type |
| pegRNA11 | -4 to -1 TCAC sub and +5 G to A | 13.33% | 6.25% | 6.67% | Wild type |
| pegRNA12 | -5 to -1 TTCAC sub and +5 G to A | 0.00% | 0.00% | 0.00% | Wild type |
| pegRNA21 | -5 to -1 GTCAC sub and +5 G to A | 6.25% | 0.00% | 0.00% | Wild type |
| pegRNA22 | -5 to -1 CTCAC sub and +5 G to A | 6.25% | 0.00% | 0.00% | Wild type |
| pegRNA13 | -6 to -1 GTTCAC sub and +5 G to A | 6.25% | 0.00% | 0.00% | Wild type |
| pegRNA14 | -7 to -1 AGTTCAC sub and +5 G to A | 0.00% | 0.00% | 0.00% | Wild type |
| pegRNA18 | -3 G to C and +5 G to A | 33.33% | 37.50% | 18.75% | Wild type |
| pegRNA19 | -7 T to A and +5 G to A | 0.00% | 0.00% | 0.00% | Wild type |
| pegRNA17 | -3 to -1 CAC sub and +5 G to A | 12.50% | 37.50% | 43.75% | Wild type |
| pegRNA15 | -5 to -1 TTCAC sub and +5 G to A | 12.50% | 0.00% | 0.00% | Wild type |
| pegRNA20 | -7 to -1 AGTTCAC sub and +5 G to A | 12.50% | 6.25% | 6.25% | Wild type |
| pegRNA04 | -3 G to C and +5 G to A | 6.25% | 6.25% | 18.75% | Wild type |
| pegRNA04 | -3 G to C and +5 G to A | 50.00% | 43.75% | 68.75% | *sbcB** |
| pegRNA04 | -3 G to C and +5 G to A | 37.50% | 18.75% | 12.50% | *sbcB*xseA* exoX** |
| pegRNA10 | -3 to -1 CAC sub and +5 G to A | 12.50% | 25.00% | 31.25% | Wild type |
| pegRNA10 | -3 to -1 CAC sub and +5 G to A | 87.50% | 93.75% | 93.75% | *scbB** |
| pegRNA10 | -3 to -1 CAC sub and +5 G to A | 100.00% | 93.33% | 100.00% | *sbcB*xseA* exoX** |
| pegRNA04 | -3 G to C and +5 G to A | 25.00% | 12.50% | 12.50% | Wild type |
| pegRNA04 | -3 G to C and +5 G to A | 13.33% | 15.38% | 6.25% | *mutL** |
| pegRNA04 | -3 G to C and +5 G to A | 6.25% | 18.75% | 12.50% | *mutS** |
| pegRNA04 | -3 G to C and +5 G to A | 6.25% | 31.25% | 26.67% | *mutL*mutS** |

| | | | | | |
|---|---|---|---|---|---|
| pegRNA10 | -3 to -1 CAC sub and +5 G to A | 23.08% | 20.00% | 18.75% | Wild type |
| pegRNA10 | -3 to -1 CAC sub and +5 G to A | 18.75% | 31.25% | 31.25% | *mutL** |
| pegRNA10 | -3 to -1 CAC sub and +5 G to A | 76.92% | 83.33% | 92.31% | *mutS** |
| pegRNA10 | -3 to -1 CAC sub and +5 G to A | 93.75% | 93.75% | 100.00% | *mutL*mutS** |
| pegRNA23 | -3 G to C | 37.50% | - | - | Wild type |
| pegRNA24 | -3 G ins | 93.33% | - | - | Wild type |
| pegRNA25 | -3 G del | 6.25% | - | - | Wild type |
| pegRNA34 | -3 to -1 ATG sub | 12.50% | 12.50% | 12.50% | Wild type |
| pegRNA35 | -3 to -1 CGG sub | 12.50% | 12.50% | 12.50% | Wild type |
| pegRNA36 | -6 CAGCT ins | 12.50% | - | - | *sbcB** |
| pegRNA37 | -6 ACAGCT ins | 43.75% | - | - | *sbcB** |
| pegRNA38 | -5 to -1 AAGTT del | 93.33% | - | - | *sbcB** |
| pegRNA39 | -6 to -1 CAAGTT del | 80.00% | - | - | *sbcB** |
| Targeting at *pta* | | | | | |
| pegRNA26 | +6 G to C | 81.25% | - | - | Wild type |
| pegRNA27 | -3 A to G and +6 G to C | 0.00% | 0.00% | 0.00% | Wild type |
| pegRNA27 | -3 A to G and +6 G to C | 25.00% | 25.00% | 25.00% | *mutS** |
| pegRNA27 | -3 A to G and +6 G to C | 14.29% | 25.00% | 33.33% | *sbcB** |
| pegRNA28 | -3 to -1 TGC sub and +6 G to C | 0.00% | 0.00% | 0.00% | Wild type |
| pegRNA28 | -3 to -1 TGC sub and +6 G to C | 0.00% | 0.00% | 0.00% | *mutS** |
| pegRNA28 | -3 to -1 TGC sub and +6 G to C | 100.00% | 100.00% | 100.00% | *sbcB** |
| Targeting at *glnA* | | | | | |
| pegRNA29 | -2 to -1 TAG sub | 6.25% | - | - | *sbcB*xseA* exoX** |
| Targeting at *gyrA* | | | | | |
| pegRNA32 | -3 to -2 AC sub | 18.75% | - | - | *sbcB*xseA* exoX** |
| pegRNA33 | -1 to +2 CTC sub | 25.00% | - | - | *sbcB*xseA* exoX** |

[a.] First position downstream of the nick as position +1 of a specific gene. Ins indicate insertions, sub indicate substitutions, and del indicate deletion.

**Supplementary Table 2.** Free energy of the RNA-DNA duplexes [a].

| pegRNA | Sequence (3' – 5') | Free energy (with 13-nt PBS) (kcal $mol^{-1}$) | Free energy (with PBS for nRTP) (kcal $mol^{-1}$) |
|---|---|---|---|
| pegRNA01 | GACUUCAGUUCAA | -11.46 | -11.46 |
| pegRNA02 | GACUUCAGUUCAG | -12.14 | -10.57 |
| pegRNA03 | GACUUCAGUUCTA | -7.39 | -9.21 |
| pegRNA04 | GACUUCAGUUGAA | -7.51 | -8.04 |
| pegRNA05 | GACUUCAGUACAA | -8.92 | -7.70 |
| pegRNA06 | GACUUCAGAUCAA | -8.62 | -6.19 |
| pegRNA07 | GACUUCACUUCAA | -5.64 | -4.67 |
| pegRNA09 | GACUUCAGUUCUG | -9.67 | -9.21 |
| pegRNA10 | GACUUCAGUUGUG | -7.62 | -8.04 |
| pegRNA11 | GACUUCAGUAGUG | -5.91 | -7.70 |
| pegRNA21 | GACUUCAGCAGUG | -5.11 | -6.19 |
| pegRNA22 | GACUUCAGGAGUG | -5.40 | -6.19 |
| pegRNA14 | GACUUCACAAGUG | -0.55 | -4.67 |

[a.] A calculation matrix is provided in Source Data.

**Supplementary Table 3.** Strains used in this study.

| Name | Features | Source |
|---|---|---|
| *E. coli* DH5α | Wild type | Takara |
| *E. coli* MG1655 | Wild type | Lab stock |
| PR19 | MG1655, pGFP | This study |
| PR20 | MG1655, early stop codon insertion in *sbcB* (Gln129*). | This study |
| PR21 | MG1655, early stop codon insertion in *sbcB* (Gln129*), *xseA* (Gln116*) and *exoX* (Trp63*). | This study |
| PR22 | MG1655, early stop codon insertion in *mutL* (Gln4*). | This study |
| PR23 | MG1655, early stop codon insertion in *mutS* (Gln138*). | This study |
| PR24 | MG1655, early stop codon insertion in *mutL* (Gln4*) and *mutS* (Gln138*). | This study |
| PR25 | MG1655, DNA substitution at the position 92 in *pta*. | This study |
| PR28 | PR21, DNA substitution at the positions from 73 to 75 in *glnA*. | This study |
| PR30 | PR21, DNA substitution at the positions from 242 and 243 in *gyrA*. | This study |
| PR31 | PR21, DNA substitution at the positions from 247 to 249 in *gyrA*. | This study |
| PR32 | PR20, DNA substitution at the positions 84 and 92 in *pta*. | This study |
| PR33 | PR20, DNA substitution at the positions from 84 to 86 and 92 in *pta*. | This study |
| PR34 | PR23, DNA substitution at the positions 84 and 92 in *pta*. | This study |
| PR35 | PR23, DNA substitution at the positions from 84 to 86 and 92 in *pta*. | This study |
| PR36 | PR19, DNA substitution at the position 346 in *gfp*. | This study |
| PR37 | PR19, DNA substitution at the positions 341 and 346 in *gfp.* | This study |
| PR38 | PR19, DNA substitution at the positions 340 and 346 in *gfp.* | This study |
| PR39 | PR19, DNA substitution at the positions 339 and 346 in *gfp.* | This study |
| PR40 | PR19, DNA substitution at the positions 338 and 346 in *gfp.* | This study |
| PR41 | PR19, DNA substitution at the positions 337 and 346 in *gfp.* | This study |
| PR42 | PR19, DNA substitution at the positions 336 and 346 in *gfp.* | This study |
| PR43 | PR19, DNA substitution at the positions 340 to 341, and 346 in *gfp.* | This study |
| PR44 | PR19, DNA substitution at the positions 339 to 341, and 346 in *gfp.* | This study |
| PR45 | PR19, DNA substitution at the positions 338 to 341, and 346 in *gfp.* | This study |
| PR46 | PR19, DNA substitution at the positions 337 to 341 (AAGTT to TTCAC), and 346 in *gfp.* | This study |

| | | |
|---|---|---|
| PR47 | PR19, DNA substitution at the positions 337 to 341 (AAGTT to GTCAC), and 346 in *gfp.* | This study |
| PR48 | PR19, DNA substitution at the positions 337 to 341 (AAGTT to CTCAC), and 346 in *gfp.* | This study |
| PR49 | PR19, DNA substitution at the positions 336 to 341, and 346 in *gfp.* | This study |
| PR50 | PR19, DNA substitution at the positions 335 to 341, and 346 in *gfp.* | This study |
| PR51 | PR19, DNA insertion at the position 339 in *gfp*. | This study |
| PR52 | PR19, DNA deletion at the position 339 in *gfp*. | This study |
| PR53 | PR19, DNA substitution at the position 339 in *gfp*. | This study |
| PR54 | PR20, pGFP | This study |
| PR55 | PR21, pGFP | This study |
| PR56 | PR22, pGFP | This study |
| PR57 | PR23, pGFP | This study |
| PR58 | PR24, pGFP | This study |
| PR59 | PR53, DNA substitution at the positions 339 and 346 in *gfp*. | This study |
| PR60 | PR53, DNA substitution at the positions 339 to 341, and 346 in *gfp.* | This study |
| PR61 | PR54, DNA substitution at the positions 339 and 346 in *gfp.* | This study |
| PR62 | PR54, DNA substitution at the positions 339 to 341, and 346 in *gfp.* | This study |
| PR63 | PR55, DNA substitution at the positions 339 and 346 in *gfp.* | This study |
| PR64 | PR55, DNA substitution at the positions 339 to 341, and 346 in *gfp.* | This study |
| PR65 | PR56, DNA substitution at the positions 339 to 341, and 346 in *gfp.* | This study |
| PR66 | PR56, DNA substitution at the positions 339 to 341, and 346 in *gfp.* | This study |
| PR67 | PR57, DNA substitution at the positions 339 to 341, and 346 in *gfp.* | This study |
| PR68 | PR57, DNA substitution at the positions 339 to 341, and 346 in *gfp.* | This study |
| PR69 | PR19, DNA substitution at the positions 111 to 113 in *gfp.* | This study |
| PR70 | PR19, DNA substitution at the positions 264 to 266 in *gfp.* | This study |
| PR71 | PR54, 5 bp DNA insertion at the positions 337. | This study |
| PR72 | PR54, 6 bp DNA insertion at the positions 337. | This study |
| PR73 | PR54, 5 bp DNA deletion at the positions 337. | This study |
| PR74 | PR54, 6 bp DNA deletion at the positions 336. | This study |

[a.] Position is counted from the first nucleotide of a specific gene.

**Supplementary Table 4.** pegRNAs sequences used in this study.

| pegRNA | Target | PAM | spacer | PBS (5'-3') | RTT (5'-3') |
|---|---|---|---|---|---|
| pegRNA01 | *gfp* | AGG | agagctgaagtcaagtttga | aacttgacttcag | ggtatcactttca |
| pegRNA02 | *gfp* | AGG | agagctgaagtcaagtttga | gacttgacttcag | ggtatcactttca |
| pegRNA03 | *gfp* | AGG | agagctgaagtcaagtttga | atcttgacttcag | ggtatcactttca |
| pegRNA04 | *gfp* | AGG | agagctgaagtcaagtttga | aagttgacttcag | ggtatcactttca |
| pegRNA05 | *gfp* | AGG | agagctgaagtcaagtttga | aacatgacttcag | ggtatcactttca |
| pegRNA06 | *gfp* | AGG | agagctgaagtcaagtttga | aactagacttcag | ggtatcactttca |
| pegRNA07 | *gfp* | AGG | agagctgaagtcaagtttga | aacttcacttcag | ggtatcactttca |
| pegRNA08 | *gfp* | AGG | agagctgaagtcaagtttga | aacttgtcttcag | ggtatcactttca |
| pegRNA09 | *gfp* | AGG | agagctgaagtcaagtttga | gtcttgacttcag | ggtatcactttca |
| pegRNA10 | *gfp* | AGG | agagctgaagtcaagtttga | gtgttgacttcag | ggtatcactttca |
| pegRNA11 | *gfp* | AGG | agagctgaagtcaagtttga | gtgatgacttcag | ggtatcactttca |
| pegRNA12 | *gfp* | AGG | agagctgaagtcaagtttga | gtgaagacttcag | ggtatcactttca |
| pegRNA13 | *gfp* | AGG | agagctgaagtcaagtttga | gtgaacacttcag | ggtatcactttca |
| pegRNA14 | *gfp* | AGG | agagctgaagtcaagtttga | gtgaactcttcag | ggtatcactttca |
| pegRNA15 | *gfp* | AGG | agagctgaagtcaagtttga | gtgaagacttcagct | ggtatcactttca |
| pegRNA17 | *gfp* | AGG | agagctgaagtcaagtttga | gtgttgacttcagct | ggtatcactttca |
| pegRNA18 | *gfp* | AGG | agagctgaagtcaagtttga | aagttgacttcagct | ggtatcactttca |
| pegRNA19 | *gfp* | AGG | agagctgaagtcaagtttga | aacttgtcttcagct | ggtatcactttca |
| pegRNA20 | *gfp* | AGG | agagctgaagtcaagtttga | gtgaactcttcagct | ggtatcactttca |
| pegRNA21 | *gfp* | AGG | agagctgaagtcaagtttga | gtgacgacttcag | ggtatcactttca |
| pegRNA22 | *gfp* | AGG | agagctgaagtcaagtttga | gtgaggacttcag | ggtatcactttca |
| pegRNA23 | *gfp* | AGG | agagctgaagtcaagtttga | aagttgacttcag | ggtatcaccttca |
| pegRNA24 | *gfp* | AGG | agagctgaagtcaagtttga | aagcttgacttcag | ggtatcaccttca |
| pegRNA25 | *gfp* | AGG | agagctgaagtcaagtttga | aa-ttgacttcag | ggtatcaccttca |
| pegRNA26 | *pta* | AGG | atccgtgcaatggaacgcaa | cgttccattgcac | acgaacggctttg |
| pegRNA27 | *pta* | AGG | atccgtgcaatggaacgcaa | cgctccattgcac | acgaacggctttg |
| pegRNA28 | *pta* | AGG | atccgtgcaatggaacgcaa | gcatccattgcac | acgaacggctttg |
| pegRNA29 | *glnA* | AGG | ttgcgcttcaccgatactaa | taatcggtgaagc | ttctttacctttc |
| pegRNA32 | *gyrA* | CGG | ccatccccatggtgactcgg | agtcaccatgggg | tcatagaccgccg |
| pegRNA33 | *gyrA* | CGG | ataccatccccatggtgact | caccatggggatg | tagaccgccgagt |
| pegRNA34 | *gfp* | CGG | ggtgaaggtgatgctactta | catgcatcacctt | caatttaccgtaa |
| pegRNA35 | *gfp* | AGG | ttcaagtctgccatgccaga | ccgatggcagact | aacataaccttct |
| pegRNA36 | *gfp* | AGG | agagctgaagtcaagtttga | aacttagctggacttcag | ggtatcaccttca |

| | | | | | |
|---|---|---|---|---|---|
| pegRNA37 | *gfp* | AGG | agagctgaagtcaagtttga | aacttagctgtgacttcag | ggtatcaccttca |
| pegRNA38 | *gfp* | AGG | agagctgaagtcaagtttga | gacttcag | ggtatcaccttca |
| pegRNA39 | *gfp* | AGG | agagctgaagtcaagtttga | acttcag | ggtatcaccttca |
| pegRNA40 | *gfp* | AGG | agagctgaagtcaagtttga | aacttgacttcag | ccaataacccactcacgttacacgttcctcggtatcactttca |
| pegRNA41 | *gfp* | AGG | agagctgaagtcaagtttga | gtgttgacttcag | ccaataacccactcacgttacacgttcctcggtatcactttca |
| pegRNA42 | *gfp* | AGG | agagctgaagtcaagtttga | gtgttgacttcag | ccaataacccactcacgttacacgttcctcggtatcaccttca |

**Supplementary Table 5.** gRNAs sequences used for base editing in this study.

| gRNA | Target | PAM | Strand [a] | spacer |
|---|---|---|---|---|
| gRNA-mutS | *mutS* | CGG | C | tggcaggacagcaaaggttt |
| gRNA-mutL | *mutL* | TGG | C | tcaggtcttaccgccacaac |
| gRNA-sbcB | *sbcB* | GGG | C | gcagcatgataactcgcgct |
| gRNA-xseA | *xseA* | AGG | C | acagctcaaagcgaagttgc |
| gRNA-exoX | *exoX* | TGG | N | ccacggtttatcggcgacca |

[a] C stands for coding strand and N stands for non-coding strand.

**Supplementary Table 6.** Plasmids used in this study.

| Name | Description | Source |
|---|---|---|
| pBR322 | *ColE1* ori, *bla*, tetA | Lab stock [1] |
| pBBR1MCS-5 | *pBBR1* ori, pBBR1 Rep, *GmR* | Lab stock [2] |
| pAM4787 | *ColE1* ori, partial sequence from pANS | Addgene #120088 [3] |
| pCMV-PE2-P2A-GFP | Cas9 (H840A)-M-MLVrt (D200N, T306K, W313F, T330P, L603W) | Addgene #132776 [4] |
| pBR322-p15A | pBR322, *p15A* ori | Lab stock |
| pBeCas9 | *oriR101*, *bla*, trc promoter, dCas9-PmCDA1-ugi | Lab stock [5] |
| pPE.S | PE.S (ncas9 from *S. pyogenes* with native sequence fused with *E. coli* codon-optimized RT), *lacI*-$P_{trc}$, *GmR*, *oriR101* | Lab stock [6] |
| pGFP | *pBBR1* ori, pBBR1 Rep, *bla*, *gfp* | This study |
| pRC | pPE.S, replace *oriR101* with *p15A* ori | This study |
| pRC-His | pRC, His-tag | This study |
| pPegRNA01 | pUC57, pegRNA01 | This study |
| pPegRNA02 | pUC57, pegRNA02 | This study |
| pPegRNA03 | pUC57, pegRNA03 | This study |
| pPegRNA04 | pUC57, pegRNA04 | This study |
| pPegRNA05 | pUC57, pegRNA05 | This study |
| pPegRNA06 | pUC57, pegRNA06 | This study |
| pPegRNA07 | pUC57, pegRNA07 | This study |
| pPegRNA08 | pUC57, pegRNA08 | This study |
| pPegRNA09 | pUC57, pegRNA09 | This study |
| pPegRNA10 | pUC57, pegRNA10 | This study |
| pPegRNA11 | pUC57, pegRNA11 | This study |
| pPegRNA12 | pUC57, pegRNA12 | This study |
| pPegRNA13 | pUC57, pegRNA13 | This study |
| pPegRNA14 | pUC57, pegRNA14 | This study |
| pPegRNA15 | pUC57, pegRNA15 | This study |
| pPegRNA17 | pUC57, pegRNA17 | This study |
| pPegRNA18 | pUC57, pegRNA18 | This study |
| pPegRNA19 | pUC57, pegRNA19 | This study |
| pPegRNA20 | pUC57, pegRNA20 | This study |
| pPegRNA21 | pUC57, pegRNA21 | This study |
| pPegRNA22 | pUC57, pegRNA22 | This study |

| | | |
|---|---|---|
| pPegRNA23 | pUC57, pegRNA23 | This study |
| pPegRNA24 | pUC57, pegRNA24 | This study |
| pPegRNA25 | pUC57, pegRNA25 | This study |
| pPegRNA26 | pUC57, pegRNA16 | This study |
| pPegRNA27 | pUC57, pegRNA27 | This study |
| pPegRNA28 | pUC57, pegRNA28 | This study |
| pPegRNA29 | pUC57, pegRNA29 | This study |
| pPegRNA32 | pUC57, pegRNA32 | This study |
| pPegRNA33 | pUC57, pegRNA33 | This study |
| pRC12 | pRC, PegRNA01 | This study |
| pRC13 | pRC, PegRNA02 | This study |
| pRC14 | pRC, PegRNA03 | This study |
| pRC15 | pRC, PegRNA04 | This study |
| pRC16 | pRC, PegRNA05 | This study |
| pRC17 | pRC, PegRNA06 | This study |
| pRC18 | pRC, PegRNA07 | This study |
| pRC19 | pRC, PegRNA08 | This study |
| pRC20 | pRC, PegRNA09 | This study |
| pRC21 | pRC, PegRNA10 | This study |
| pRC22 | pRC, PegRNA11 | This study |
| pRC23 | pRC, PegRNA12 | This study |
| pRC24 | pRC, PegRNA13 | This study |
| pRC25 | pRC, PegRNA14 | This study |
| pRC26 | pRC, PegRNA15 | This study |
| pRC27 | pRC, PegRNA17 | This study |
| pRC28 | pRC, PegRNA18 | This study |
| pRC29 | pRC, PegRNA19 | This study |
| pRC30 | pRC, PegRNA20 | This study |
| pRC31 | pRC, PegRNA21 | This study |
| pRC32 | pRC, PegRNA22 | This study |
| pRC33 | pRC, PegRNA23 | This study |
| pRC34 | pRC, PegRNA24 | This study |
| pRC35 | pRC, PegRNA25 | This study |
| pRC36 | pRC, PegRNA26 | This study |
| pRC37 | pRC, PegRNA27 | This study |
| pRC38 | pRC, PegRNA28 | This study |
| pRC39 | pRC, PegRNA29 | This study |
| pRC40 | pRC, PegRNA32 | This study |

| | | |
|---|---|---|
| pRC41 | pRC, PegRNA33 | This study |
| pRC42 | pRC, PegRNA34 | This study |
| pRC43 | pRC, PegRNA35 | This study |
| pRC44 | pRC, PegRNA36 | This study |
| pRC45 | pRC, PegRNA37 | This study |
| pRC46 | pRC, PegRNA38 | This study |
| pRC47 | pRC, PegRNA39 | This study |
| pBeCas9-mutS | pBeCas9, gRNA-mutS | This study |
| pBeCas9-mutL | pBeCas9, gRNA-mutL | This study |
| pBeCas9-sbcB | pBeCas9, gRNA-sbcB | This study |
| pBeCas9-xseA | pBeCas9, gRNA-xseA | This study |
| pBeCas9-exoX | pBeCas9, gRNA-exoX | This study |

**Supplementary Table 7.** Primers used in this study.

| Primer | Sequence |
|---|---|
| **Primers for In-Fusion DNA assembly** | |
| XIA-PRC-113 | CATTTGAGAAGCACACGGTCACAGGAAACAGCTATGACCG |
| XIA-PRC-114 | TTGACTACCGGAAGCAGTGTTCTAGATTGTAAAACGACGGCCAGTC |
| XIA-PRC-132 | CACACAGGAAACAGACCATGATGGATAAGAAATACTCAATAGG |
| XIA-PRC-135 | CCTATTGAGTATTTCTTATCCATCATGGTCTGTTTCCTGTGTG |
| XIA-PRC-276 | AATAGGCGTATCACGAGGCCAGGCATTTCTGTCCTGGCTG |
| XIA-PRC-277 | GTAGGTGTTCCACAGGGTAGTGCCCAGCTCGGTCTAGAAT |
| XIA-PRC-278 | ATTCTAGACCGAGCTGGGCACTACCCTGTGGAACACCTAC |
| XIA-PRC-279 | CAGCCAGGACAGAAATGCCTGGCCTCGTGATACGCCTATT |
| XIA-PRC-283 | AGGATCTAGGTGAAGATCCTCTTTGCTTCGCAAAGTCGTG |
| XIA-PRC-284 | CGTATCGTGAGCATCCTCTCGGGAGGCAGACAAGGTATAG |
| XIA-PRC-285 | CACGACTTTGCGAAGCAAAGAGGATCTTCACCTAGATCCT |
| XIA-PRC-286 | CTATACCTTGTCTGCCTCCCGAGAGGATGCTCACGATACG |
| XIA-PRC-291 | TGGCAGGACAGCAAAGGTTTGTTTTAGAGCTAGAAATAGC |
| XIA-PRC-292 | AAACCTTTGCTGTCCTGCCAGCTAGCATTATACCTAGGAC |
| XIA-PRC-298 | GACTGGCCGTCGTTTTACAATCTAGAACACTGCTTCCGGTAGTCAA |
| XIA-PRC-299 | CGGTCATAGCTGTTTCCTGTGACCGTGTGCTTCTCAAATG |
| XIA-PRC-303 | TCAGGTCTTACCGCCACAACGTTTTAGAGCTAGAAATAGC |
| XIA-PRC-304 | GTTGTGGCGGTAAGACCTGAGCTAGCATTATACCTAGGAC |
| XIA-PRC-310 | CGACGGTATCGATAAGCTTGATGCGAGGCATATTTATGGTGA |
| XIA-PRC-311 | CGCTCTAGAACTAGTGGATCTGTTACTTGGTTCTGGCGAG |
| XIA-PRC-312 | TCACCATAAATATGCCTCGCATCAAGCTTATCGATACCGTCG |
| XIA-PRC-313 | CTCGCCAGAACCAAGTAACAGATCCACTAGTTCTAGAGCG |
| XIA-PRC-314 | GCGGATTTGAACGTTGCGAATCCTTGACAGCTAGCTCAGT |
| XIA-PRC-315 | CCAGCTCGGTCTAGATTGCTCCTTTGAGTGAGCTGATACC |
| XIA-PRC-316 | ACTGAGCTAGCTGTCAAGGATTCGCAACGTTCAAATCCGC |
| XIA-PRC-317 | GGTATCAGCTCACTCAAAGGAGCAATCTAGACCGAGCTGG |
| XIA-PRC-387 | CCGCAGGAAGCACGGGCGAAGTTTTAGAGCTAGAAATAGC |
| XIA-PRC-388 | TTCGCCCGTGCTTCCTGCGGGCTAGCATTATACCTAGGAC |
| XIA-PRC-423 | ACAGCTCAAAGCGAAGTTGCGTTTTAGAGCTAGAAATAGC |
| XIA-PRC-424 | GCAACTTCGCTTTGAGCTGTGCTAGCATTATACCTAGGAC |
| XIA-PRC-427 | CCACGGTTTATCGGCGACCAGTTTTAGAGCTAGAAATAGC |
| XIA-PRC-428 | TGGTCGCCGATAAACCGTGGGCTAGCATTATACCTAGGAC |
| XIA-PRC-547 | CTCGAGCACCACCACCACCACCACTGACCATGCGAGAGTAGGGAACT |
| XIA-PRC-548 | TCAGTGGTGGTGGTGGTGGTGCTCGAGTTCGAATTCAGAACCGTCAG |

| | |
|---|---|
| XIA-PRC-549 | GTAAAACGACGGCCAGTAAG |
| XIA-PRC-550 | GTAAAACGACGGCCAGTAAG |
| **Primers for colony PCR and sequencing** | |
| XIA-PRC-287 | GTTCCATGGCCAACCTTAGT |
| XIA-PRC-288 | GTTCCAGACTATCGGCTGTA |
| XIA-PRC-290 | TGGACCATCACCAATTGGAG |
| XIA-PRC-293 | CTCTGGTGATTCAGGACTCT |
| XIA-PRC-294 | CCTACCTACGTAACGGACTA |
| XIA-PRC-295 | ATGCAGCAGTATCTCAGGCT |
| XIA-PRC-296 | CGACGGCCTTCAATTAACGA |
| XIA-PRC-297 | AACTGGTGAATCAGGGAGAG |
| XIA-PRC-300 | CAATCGAGGAGCCGTCAAAC |
| XIA-PRC-301 | GAACGTACCGGATTGTTGGA |
| XIA-PRC-302 | CCACCTTCATATTGGGTGGA |
| XIA-PRC-305 | GTAGGGTATGATGTAGCGAC |
| XIA-PRC-306 | GGGTGTTGTAGAACAGATCC |
| XIA-PRC-307 | CGACGATTACCAACAACAGC |
| XIA-PRC-308 | CCCTGCTGATCGAAAACTCT |
| XIA-PRC-309 | ATGTTGTTGCTCAGGTCGCA |
| XIA-PRC-392 | CTCTGGCAGACAGCAGAAAT |
| XIA-PRC-393 | TGTCGCTATCCAGTTCCAGT |
| XIA-PRC-394 | GGTTTCATCTGCGGAACATC |
| XIA-PRC-429 | AATAACCAGCGCAGATAGCC |
| XIA-PRC-430 | TTGCCGTGCTGATGGCAAAT |
| XIA-PRC-431 | GATATCGGCCATCTGTTCTG |
| XIA-PRC-432 | TTCGTCTGCTGCTTGAGCAT |
| XIA-PRC-433 | AACCGGAATGCGGCTGGTAA |
| XIA-PRC-434 | CACAGCATGGGCAACAAGTT |
| XIA-PRC-497 | GATAGTGAAAGTGAACGCGG |
| XIA-PRC-498 | TAACTGGCCCACTTTGTCGA |
| XIA-PRC-499 | TTCCGGCACTGTTTAACGGT |
| XIA-PRC-500 | GAACATCTTCCAGCAACACG |
| XIA-PRC-501 | TGGTGCGCATGATAACGCCT |
| XIA-PRC-502 | CAGTGGAACGCAGGTAATCT |
| XIA-PRC-503 | TCAAGGATGTCGCAACGGAT |
| XIA-PRC-504 | ATAGCGGTTAGATGAGCGAC |
| XIA-PRC-505 | TTGCCATACCTACGGCGATA |
| XIA-PRC-506 | TGCGATGTCGGTCATTGTTG |

| | |
|---|---|
| XIA-PRC-533 | TAATGGTACCCAGATCTCCG |
| XIA-PRC-551 | GCTGACCTCTGAAGGTAAAG |
| XIA-PRC-552 | CGCTAACGGATTCACCGTTT |
| XIA-PRC-553 | TAACCGTATGGCTGACCAGG |
| **Primers for PCR and sequencing in *in vitro* experiment** | |
| XIA-PRC-554 | TGGCCAACCTTAGTCACTAC |
| XIA-PRC-555 | ACCCACTCACGTTACACGTT |
| XIA-PRC-556 | TTCAAGTCTGCCATGCCAGA |
| XIA-PRC-557 | TTTGTTTGTCAGCCATGATG |

**Supplementary Table 8.** Sequences of the target genes used in this study.

| Gene | Sequence [a, b] |
|---|---|
| *gfp* | ATGTCTAAAGGTGAAGAATTATTCACTGGTGTTGTCCCAATTTTGGTTGAATT<br>AGATGGTGATGTTAATGGTCACAAATTTTCTGTCTCCGGTGAAGGTGAAGGT<br>GATGCTACTTACGGTAAATTGACCTTAAAATTTATTTGTACTACTGGTAAATTG<br>CCAGTTCCATGGCCAACCTTAGTCACTACTTTAACTTATGGTGTTCAATGTTTT<br>TCTAGATACCCAGATCATATGAAACAACATGACTTTTTCAAGTCTGCCATGCC<br>AGAAGGTTATGTTCAAGAAAGAACTATTTTTTTCAAAGATGACGGTAACTACA<br>AGACCAGAGCTGAAGTCAAGTTTGAAGGTGATACCTTAGTTAATAGAATCGAA<br>TTAAAAGGTATTGATTTTAAAGAAGATGGTAACATTTTAGGTCACAAATTGGAA<br>TACAACTATAACTCTCACAATGTTTACATCATGGCTGACAAACAAAAGAATGG<br>TATCAAAGTTAACTTCAAAATTAGACACAACATTGAAGATGGTTCTGTTCAATT<br>AGCTGACCATTATCAACAAAATACTCCAATTGGTGATGGTCCAGTCTTGTTAC<br>CAGACAACCATTACTTATCCACTCAATCTAAATTATCCAAAGATCCAAACGAA<br>AAGAGAGACCACATGGTCTTGTTAGAATTTGTTACTGCTGCTGGTATTACCCA<br>TGGTATGGATGAATTGTACAAATAA |
| *pta*<br>(ECK2291) | GTGTCCCGTATTATTATGCTGATCCCTACCGGAACCAGCGTCGGTCTGACCA<br>GCGTCAGCCTTGGCGTGATCCGTGCAATGGAACGCAAAGGCGTTCGTCTGA<br>GCGTTTTCAAACCTATCGCTCAGCCGCGTACCGGTGGCGATGCGCCCGATC<br>AGACTACGACTATCGTGCGTGCGAACTCTTCCACCACGACGGCCGCTGAAC<br>CGCTGAAAATGAGCTACGTTGAAGGTCTGCTTTCCAGCAATCAGAAAGATGT<br>GCTGATGGAAGAGATCGTCGCAAACTACCACGCTAACACCAAAGACGCTGAA<br>GTCGTTCTGGTTGAAGGTCTGGTCCCGACACGTAAGCACCAGTTTGCCCAGT<br>CTCTGAACTACGAAATCGCTAAAACGCTGAATGCGGAAATCGTCTTCGTTATG<br>TCTCAGGGCACTGACACCCCGGAACAGCTGAAAGAGCGTATCGAACTGACC<br>CGCAACAGCTTCGGCGGTGCCAAAAACACCAACATCACCGGCGTTATCGTTA<br>ACAAACTGAACGCACCGGTTGATGAACAGGGTCGTACTCGCCCGGATCTGT<br>CCGAGATTTTCGACGACTCTTCCAAAGCTAAAGTAAACAATGTTGATCCGGC<br>GAAGCTGCAAGAATCCAGCCCGCTGCCGGTTCTCGGCGCTGTGCCGTGGAG<br>CTTTGACCTGATCGCGACTCGTGCGATCGATATGGCTCGCCACCTGAATGCG<br>ACCATCATCAACGAAGGCGACATCAATACTCGCCGCGTTAAATCCGTCACTT<br>TCTGCGCACGCAGCATTCCGCACATGCTGGAGCACTTCCGTGCCGGTTCTCT<br>GCTGGTGACTTCCGCAGACCGTCCTGACGTGCTGGTGGCCGCTTGCCTGGC<br>AGCCATGAACGGCGTAGAAATCGGTGCCCTGCTGCTGACTGGCGGTTACGA<br>AATGGACGCGCGCATTTCTAAACTGTGCGAACGTGCTTTCGCTACCGGCCTG<br>CCGGTATTTATGGTGAACACCAACACCTGGCAGACCTCTCTGAGCCTGCAGA<br>GCTTCAACCTGGAAGTTCCGGTTGACGATCACGAACGTATCGAGAAAGTTCA |

GGAATACGTTGCTAACTACATCAACGCTGACTGGATCGAATCTCTGACTGCC
ACTTCTGAGCGCAGCCGTCGTCTGTCTCCGCCTGCGTTCCGTTATCAGCTGA
CTGAACT

TGCGCGCAAAGCGGGCAAACGTATCGTACTGCCGGAAGGTGACGAACCGCG
TACCGTTAAAGCAGCCGCTATCTGTGCTGAACGTGGTATCGCAACTTGCGTA
CTGCTGGGTAATCCGGCAGAGATCAACCGTGTTGCAGCGTCTCAGGGTGTA
GAACTGGGTGCAGGGATTGAAATCGTTGATCCAGAAGTGGTTCGCGAAAGCT
ATGTTGGTCGTCTGGTCGAACTGCGTAAGAACAAAGGCATGACCGAAACCGT
TGCCCGCGAACAGCTGGAAGACAACGTGGTGCTCGGTACGCTGATGCTGGA
ACAGGATGAAGTTGATGGTCTGGTTTCCGGTGCTGTTCACACTACCGCAAAC
ACCATCCGTCCGCCGCTGCAGCTGATCAAAACTGCACCGGGCAGCTCCCTG
GTATCTTCCGTGTTCTTCATGCTGCTGCCGGAACAGGTTTACGTTTACGGTG
ACTGTGCGATCAACCCGGATCCGACCGCTGAACAGCTGGCAGAAATCGCGA
TTCAGTCCGCTGATTCCGCTGCGGCCTTCGGTATCGAACCGCGCGTTGCTAT
GCTCTCCTACTCCACCGGTACTTCTGGTGCAGGTAGCGACGTAGAAAAAGTT
CGCGAAGCAACTCGTCTGGCGCAGGAAAAACGTCCTGACCTGATGATCGAC
GGTCCGCTGCAGTACGACGCTGCGGTAATGGCTGACGTTGCGAAATCCAAA
GCGCCGAACTCTCCGGTTGCAGGTCGCGCTACCGTGTTCATCTTCCCGGAT
CTGAACACCGGTAACACCACCTACAAAGCGGTACAGCGTTCTGCCGACCTGA
TCTCCATCGGGCCGATGCTGCAGGGTATGCGCAAGCCGGTTAACGACCTGT
CCCGTGGCGCACTGGTTGACGATATCGTCTACACCATCGCGCTGACTGCGA
TTCAGTCTGCACAGCAGCAGTAA

ATGAGTGCAATAGAAAATTTCGACGCCCATACGCCCATGATGCAGCAGTATC
TCAGGCTGAAAGCCCAGCATCCCGAGATCCTGCTGTTTTACCGGATGGGTGA
TTTTTATGAACTGTTTTATGACGACGCAAAACGCGCGTCGCAACTGCTGGATA
TTTCACTGACCAAACGCGGTGCTTCGGCGGGAGAGCCGATCCCGATGGCGG
GGATTCCCTACCATGCGGTGGAAAACTATCTCGCCAAACTGGTGAATCAGGG
AGAGTCCGTTGCCATCTGCGAACAAATTGGCGATCCGGCGACCAGCAAAGG
TCCGGTTGAGCGCAAAGTTGTGCGTATCGTTACGCCAGGCACCATCAGCGAT

*mutS* (ECK2728)

GAAGCCCTGTTGCAGGAGCGTCAGGACAACCTGCTGGCGGCTATCTGGCAG
GACAGCAAAGGTTTCGGCTACGCGACGCTGGATATCAGTTCCGGGCGTTTTC
GCCTGAGCGAACCGGCTGACCGCGAAACGATGGCGGCAGAACTGCAACGC
ACTAATCCTGCGGAACTGCTGTATGCAGAAGATTTTGCTGAAATGTCGTTAAT
TGAAGGCCGTCGCGGCCTGCGCCGTCGCCCGCTGTGGGAGTTTGAAATCGA
CACCGCGCGCCAGCAGTTGAATCTGCAATTTGGGACCCGCGATCTGGTCGG
TTTTGGCGTCGAGAACGCGCCGCGCGGACTTTGTGCTGCCGGTTGTCTGTT
GCAGTATGCGAAAGATACCCAACGTACGACTCTGCCGCATATTCGTTCCATC
ACCATGGAACGTGAGCAGGACAGCATCATTATGGATGCCGCGACGCGTCGT

AATCTGGAAATCACCCAGAACCTGGCGGGTGGTGCGGAAAATACGCTGGCT
TCTGTGCTCGACTGCACCGTCACGCCGATGGGCAGCCGTATGCTGAAACGC
TGGCTGCATATGCCAGTGCGCGATACCCGCGTGTTGCTTGAGCGCCAGCAA
ACTATTGGCGCATTGCAGGATTTCACCGCCGGGCTACAGCCGGTACTGCGT
CAGGTCGGCGACCTGGAACGTATTCTGGCACGTCTGGCTTTACGAACTGCTC
GCCCACGCG

ATCTGGCCCGTATGCGCCACGCTTTCCAGCAACTGCCGGAGCTGCGTGCGC
AGTTAGAAACTGTCGATAGTGCACCGGTACAGGCGCTACGTGAGAAGATGG
GCGAGTTTGCCGAGCTGCGCGATCTGCTGGAGCGAGCAATCATCGACACAC
CGCCGGTGCTGGTACGCGACGGTGGTGTTATCGCATCGGGCTATAACGAAG
AGCTGGATGAGTGGCGCGCGCTGGCTGACGGCGCGACCGATTATCTGGAG
CGTCTGGAAGTCCGCGAGCGTGAACGTACCGGCCTGGACACGCTGAAAGTT
GGCTTTAATGCGGTGCACGGCTACTACATTCAAATCAGCCGTGGGCAAAGCC
ATCTGGCACCCATCAACTACATGCGTCGCCAGACGCTGAAAAACGCCGAGC
GCTACATCATTCCAGAGCTAAAAGAGTACGAAGATAAAGTTCTCACCTCAAAA
GGCAAAGCACTGGCACTGGAAAAACAGCTTTATGAAGAGCTGTTCGACCTGC
TGTTGCCGCATCTGGAAGCGTTGCAACAGAGCGCGAGCGCGCTGGCGGAAC
TCGACGTGCTGGTTAACCTGGCGGAACGGGCCTATACCCTGAACTACACCT
GCCCGACCTTCATTGATAAACCGGGCATTCGCATTACCGAAGGTCGCCATCC
GGTAGTTGAACAAGTACTGAATGAGCCATTTATCGCCAACCCGCTGAATCTG
TCGCCGCAGCGCCGCATGTTGATCATCACCGGTCCGAACATGGGCGGTAAA
AGTACCTATATGCGCCAGACCGCACTGATTGCGCTGATGGCCTACATCGGCA
GCTATGTACCGGCACAAAAAGTCGAGATTGGACCTATCGATCGCATCTTTAC
CCGCGTAGGCGCGGCAGATGACCTGGCGTCCGGGCGCTCAACCTTTATGGT
GGAGATGACTGAAACCGCCAATATTTTACATAACGCCACCGAATACAGTCTG
GTGTTAATGGATGAGATCGGGCGTGGAACGTCCACCTACGATGGTCTGTCG
CTGGCGTGGGCGTGCGCGGAAAATCTGGCGAATAAGATTAAGGCATTGACG
TTATTTGCTACCCACTATTTCGAGCTGACCCAGTTACCGGAGAAAATGGAAG
GCGTCGCTAACGTGCATCTCGATGCACTGGAGCACGGCGACACCATTGCCT
TTATGCACAGCGTGCAGGATGGCGCGGCGAGCAAAAGCTACGGCCTGGCG
GTTGCAGCTCTGGCAGGCGTGCCAAAAGAGGTTATTAAGCGCGCACGGCAA
AAGCTGCGTGAGCTGGAAAGCATTTCGCCGAACGCCGCCGCTACGCAAGTG
GATGGTACGCAAATGTCTTTGCTGTCAGTACCAGAAGAAACTTCGCCTGCGG
TCGAAGCTCTGGAAAATCTTGATCCGGATTCACTCACCCCGCGTCAGGCGCT
GGAGTGGATTTATCGCTTGAAGAGCCTGGTGTAA

| | |
|---|---|
| *mutL* (ECK4166) | ATGCCAATTCAGGTCTTACCGCCACAACTGGCGAACCAGATTGCCGCAGGTG<br>AGGTGGTCGAGCGACCTGCGTCGGTAGTCAAAGAACTAGTGGAAAACAGCC<br>TCGATGCAGGTGCGACGCGTATCGATATTGATATCGAACGCGGTGGGGCGA |

AACTTATCCGCATTCGTGATAACGGCTGCGGTATCAAAAAAGATGAGCTGGC
GCTGGCGCTGGCTCGTCATGCCACCAGTAAAATCGCCTCTCTGGACGATCTC
GAAGCCATTATCAGCCTGGGCTTTCGCGGTGAGGCGCTGGCGAGTATCAGT
TCGGTTTCCCGCCTGACGCTCACTTCACGCACCGCAGAACAGCAGGAAGCC
TGGCAGGCCTATGCCGAAGGGCGCGATATGAACGTGACGGTAAAACCGGCG
GCGCATCCTGTGGGGACGACGCTGGAGGTGCTGGATCTGTTCTACAACACC
CCGGCGCGGCGCAAATTCCTGCGCACCGAGAAAACCGAATTTAACCACATT
GATGAGATCATCCGCCGCATTGCGCTGGCGCGTTTCGACGTCACGATCAAC
CTGTC

GCATAACGGTAAAATTGTGCGTCAGTACCGCGCAGTGCCGGAAGGCGGGCA
AAAAGAACGGCGCTTAGGCGCGATTTGCGGCACCGCTTTTCTTGAACAAGCG
CTGGCGATTGAATGGCAACACGGCGATCTCACGCTACGCGGCTGGGTGGCC
GATCCAAATCACACCACGCCCGCACTGGCAGAAATTCAGTATTGCTACGTGA
ACGGTCGCATGATGCGCGATCGCCTGATCAATCACGCGATCCGCCAGGCCT
GCGAAGACAAACTGGGGGCCGATCAGCAACCGGCATTTGTGTTGTATCTGG
AGATCGACCCACATCAGGTGGACGTCAACGTGCACCCCGCCAAACACGAAG
TGCGTTTCCATCAGTCGCGTCTGGTGCATGATTTTATCTATCAGGGCGTGCT
GAGCGTGCTACAACAGCAACTGGAAACGCCGCTACCGCTGGACGATGAACC
CCAACCTGCACCGCGTTCCATTCCGGAAAACCGCGTGGCGGCGGGGCGCA
ATCACTTTGCAGAACCGGCAGCTCGTGAGCCGGTAGCTCCGCGCTACACTC
CTGCGCCAGCATCAGGCAGTCGTCCGGCTGCCCCCTGGCCGAATGCGCAG
CCAGGCTACCAGAAACAGCAAGGTGAAGTGTATCGCCAGCTTTTGCAAACGC
CCGCGCCGATGCAAAAATTAAAAGCGCCGGAACCGCAGGAACCTGCACTTG
CGGCGAACAGTCAGAGTTTTGGTCGGGTACTGACTATCGTCCATTCCGACTG
TGCGTTGCTGGAGCGCGACGGCAACATTTCACTTTTATCCTTGCCAGTGGCA
GAACGTTGGCTGCGTCAGGCACAATTGACGCCGGGTGAAGCGCCCGTTTGC
GCCCAGCCGCTGCTGATTCCGTTGCGGCTAAAAGTTTCTGCCGAAGAAAAAT
CGGCATTAGAAAAAGCGCAGTCTGCCCTGGCGGAATTGGGTATTGATTTCCA
GTCAGATGCACAGCATGTGACCATCAGGGCAGTGCCTTTACCCTTACGCCAA
CAAAATTTACAAATCTTGATTCCTGAACTGATAGGCTACCTGGCGAAGCAGTC
CGTATTCGAACCTGGCAATATTGCGCAGTGGATTGCACGAAATCTGATGAGC
GAACATGCGCAGTGGTCAATGGCACAGGCCATAACCCTGCTGGCGGACGTG
GAACGGTTATGTCCGCAACTTGTGAAAACGCCGCCGGGTGGTCTGTTACAAT
CTGTTGATTTACATCCGGCGATAAAAGCCCTGAAAGATGAGTGA

| | |
|---|---|
| sbcB (ECK2005) | ATGATGAATGACGGTAAGCAACAATCTACCTTTTTGTTTCACGATTACGAAAC CTTTGGCACGCACCCCGCGTTAGATCGCCCTGCACAGTTCGCAGCCATTCG CACCGATAGCGAATTCAATGTCATCGGCGAACCCGAAGTCTTTTACTGCAAG CCCGCTGATGACTATTTACCCCAGCCAGGAGCCGTATTAATTACCGGTATTA |

CCCCGCAGGAAGCACGGGCGAAAGGAGAAAACGAAGCCGCGTTTGCCGCC
CGTATTCACTCGCTTTTTACCGTACCGAAGACCTGTATTCTGGGCTACAACAA
TGTGCGTTTCGACGACGAAGTCACACGCAACATTTTTTATCGTAATTTCTACG
ATCCTTACGCCTGGAGCTGGCAGCATGATAACTCGCGCTGGGATTTACTGGA
TGTTATGCGTGCCTGTTATGCCCTGCGCCCGGAAGGAATAAACTGGCCTGAA
AATGATGACGGTCTACCGAGCTTTCGCCTTGAGCATTTAACCAAAGCGAATG
GTATTGAACATAGCAACGCCCACGATGCGATGGCTGATGTGTACGCCACTAT
TGCGATGGCAAAGCTGGTAAAAACGCGTCAGCCACGCCTGTTTGATTATCTC
TTTACCCATCGTAATAAACACAAACTGATGGCGTTGATTGATGTTCCGCAGAT
GAAACCCCTGGTGCACGTTTCCGGAATGTTTGGAGCATGGCGCGGCAATAC
CAGCTGGGTGGCACCGCTGGCGTGGCATCCTGAAAATCGCAATGCCGTAAT
TATGGTGGATTTGGCAGGAGACATTTCGCCATTACTGGAACTGGATAGCGAC
ACATTGCGCGAGCGTTTATATACCGCAAAAACCGATCTTGGCGATAACGCCG
CCGTTCCGGTTAAGCTGGTGCATATCAATAAATGTCCGGTGCTGGCCCAGGC
GAATACGCTACGCCCGGAAGATGCCGACCGACTGGGAATTAATCGTCAGCA
TTGCCTCGATAACCTGAAAATTCTGCGTGAAAATCCGCAAGTGCGCGAAAAA
GTGGTGGCGATATTCGCGGAAGCCGAACCGTTTACGCCTTCAGATAACGTG
GATGCACAGCTTTATAACGGCTTTTTCAGTGACGCAGATCGTGCAGCAATGA
AAATTGTGCTGGAAACCGAGCCGCGTAATTTACCGGCACTGGATATCACTTT
TGTTGATAAACGGATTGAAAAGCTGTTGTTCAATTATCGGGCACGCAACTTCC
CGGGGACGCTGGATTATGCCGAGCAGCAACGCTGGCTGGAGCACCGTCGC
CAGGTCTTCACGCCAGAGTTTTTGCAGGGTTATGCTGATGAATTGCAGATGC
TGGTACAACAATATGCCGATGACAAAGAGAAAGTGGCGCTGTTAAAAGCACT
TTGGCAGTACGCGGAAGAGATTGTCTAA

ATGTTACCTTCTCAATCCCCTGCAATTTTTACCGTTAGTCGCCTGAATCAAAC
GGTTCGTCTGCTGCTTGAGCATGAGATGGGACAGGTTTGGATCAGCGGCGA
AATTTCTAATTTCACGCAACCAGCTTCCGGTCACTGGTACTTTACACTCAAAG
ACGACACCGCCCAGGTACGCTGCGCGATGTTCCGCAACAGCAACCGCCGG
GTGACCTTCCGCCCACAGCATGGGCAACAAGTTTTAGTTCGCGCCAATATTA
CGCTCTACGAGCCGCGCGGCGACTACCAGATAATCGTTGAGAGTATGCAGC
xseA CGGCCGGTGAAGGGCTGCTGCAACAGAAGTACGAACAGCTCAAAGCGAAGT
(ECK2505) TGCAGGCTGAAGGTTTGTTCGATCAGCAATACAAAAAACCACTTCCCTCCCC
TGCGCATTGCGTTGGTGTGATCACCTCAAAAACCGGTGCTGCGCTACATGAT
ATTTTGCATGTGTTAAAACGTCGCGATCCTTCTCTGCCGGTGATCATCTACCC
TGCCGCCGTTCAGGGCGATGACGCGCCGGGGCAAATTGTTCGCGCCATTGA
ACTGGCGAATCAGCGCAATGAGTGCGACGTATTGATCGTCGGGCGCGGCGG
CGGTTCGCTGGAAGATTTATGGAGTTTTAACGACGAACGCGTAGCGCGGGC
GATTTTTACCAGCCGCATTCCGGTTGTCAGCGCCGTCGGGCATGAGACGGA
TGTGACCATTGCCGATTTTGTTGCCGATCTGCGTGCGCCAACGCCGTCTGCC

GCCGCTGAAGTAGTGAGCCGTAATCAGCAAGAGTTACTGCGCCAGGTGCAA
TCGACCCGTCAACGGCTGGAGATGGCGATGGATTATTATCTCGCCAACCGCA
CACGTCGCTTTACGCAAATTCATCACCGATTACAGCAACAGCATCCGCAGCT
CCGGCTGGCACGCCAGCAAACCATGCTTGAGCGCCTGCAAAAGCGAATGAG
CTTTGCGCTGGAAAATCAACTTAAGCGTACCGGGCAACAGCAGCAGCGGTTA
ACACAGCGGCTGAATCAGCAAAATCCACAGCCGAAGATTCATCGCGCGCAAA
CGCGCATTCAGCAACTGGAATATCGTTTAGCAGAAACCCTGCGCGCACAGCT
TAGCGCCACGCGTGAACGTTTCGGTAATGCAGTAACGCACCTCGAAGCCGT
AAGCCCACTGTCAACGCTGGCGCGTGGATACAGCGTTACTACTGCTACTGAC
GGCAATGTACTGAAAAAAGTGAAGCAAGTTAAAGCGGGTGAAATGCTAACCA
CACGTCTGGAAGACGGCTGGATAGAAAGTGAAGTAAAAAACATCCAGCCAGT
AAAAAAATCGCGTAAAAAGGTGCATTAA

*exoX* (ECK1845)

ATGTTGCGCATTATCGATACAGAAACCTGCGGTTTGCAGGGAGGGATCGTTG
AGATTGCCTCTGTTGATGTCATTGACGGAAAAATCGTCAACCCCATGAGCCA
CCTGGTGCGCCCCGATCGTCCTATTAGTCCACAAGCGATGGCGATTCATCGC
ATCACCGAAGCCATGGTCGCCGATAAACCGTGGATTGAAGATGTGATCCCAC
ACTATTACGGTAGTGAATGGTATGTCGCGCATAACGCCAGCTTTGACCGCCG
CGTACTGCCTGAGATGCCCGGTGAGTGGATTTGCACTATGAAACTGGCCCGT
CGTTTGTGGCCTGGGATCAAGTACAGCAATATGGCGTTATATAAAACACGCA
AGCTCAATGTACAGACGCCGCCGGGCCTGCATCATCACCGCGCGTTGTATG
ACTGTTATATCACCGCCGCGTTGCTTATCGATATTATGAACACCTCCGGCTG
GACGGCAGAACAGATGGCCGATATCACCGGACGTCCGTCGTTGATGACGAC
CTTCACCTTTGGCAAATACCGTGGCAAAGCGGTTTCCGACGTTGCCGAACGC
GATCCGGGCTATCTGCGCTGGTTATTTAATAACCTGGACAGCATGAGCCCGG
AGCTGCGTTTAACACTGAAACATTATCTGGAAAATACTTAG

*glnA* (ECK3863)

ATGTCCGCTGAACACGTACTGACGATGCTGAACGAGCACGAAGTGAAGTTTG
TTGATTTGCGCTTCACCGATACTAAAGGTAAAGAACAGCACGTCACTATCCCT
GCTCATCAGGTGAATGCTGAATTCTTCGAAGAAGGCAAAATGTTTGACGGCT
CCTCGATTGGCGGCTGGAAAGGCATTAACGAGTCCGACATGGTGCTGATGC
CAGACGCATCCACCGCAGTGATTGACCCGTTCTTCGCCGACTCCACCCTGAT
TATCCGTTGCGACATCCTTGAACCTGGCACCCTGCAAGGCTATGACCGTGAC
CCGCGCTCCATTGCGAAGCGCGCCGAAGATTACCTGCGTTCCACTGGCATT
GCCGACACCGTACTGTTCGGGCCAGAACCTGAATTCTTCCTGTTCGATGACA
TCCGTTTCGGATCATCTATCTCCGGTTCCCACGTTGCTATCGACGATATCGAA
GGCGCATGGAACTCCTCCACCCAATACGAAGGTGGTAACAAAGGTCACCGT
CCGGCAGTGAAAGGCGGTTACTTCCCGGTTCCACCGGTAGACTCGGCTCAG
GATATTCGTTCTGAAATGTGTCTGGTGATGGAACAGATGGGTCTGGTGGTTG
AAGCCCATCACCACGAAGTAGCGACTGCTGGTCAGAACGAAGTGGCTACCC

GCTTCAATACCATGACCAAAAAAGCTGACGAAATTCAGATCTACAAATATGTT
GTGCACAACGTAGCGCACCGCTTCGGTAAAACCGCGACCTTTATGCCAAAAC
CGATGTTCGGTGATAACGGCTCCGGTATGCACTGCCACATGTCTCTGTCTAA
AAACGGCGTTAACCTGTTCGCAGGCGACAAATACGCAGGTCTGTCTGAGCA
GGCGCTGTACTACATTGGCGGCGTAATCAAACACGCTAAAGCGATTAACGCC
CTGGCAAACCCGACCACCAACTCTTATAAGCGTCTGGTCCCGGGCTATGAAG
CACCGGTAATGCTGGCTTACTCTGCGCGTAACCGTTCTGCGTCTATCCGTAT
TCCGGTGGTTTCTTCTCCGAAAGCACGTCGTATCGAAGTACGTTTCCCGGAT
CCGGCAGCTAACCCGTACCTGTGCTTTGCTGCCCTGCTGATGGCCGGTCTT
GATGGTATCAAGAACAAGATCCATCCGGGCGAAGCCATGGACAAAAACCTGT
ATGACCTGCCGCCAGAAGAAGCGAAAGAGATCCCACAGGTTGCAGGCTCTC
TGGAAGAAGCACTGAACGAACTGGATCTGGACCGCGAGTTCCTGAAAGCCG
GTGGCGTGTTCACTGACGAAGCAATTGATGCGTACATCGCTCTGCGTCGCGA
AGAAGATGACCGCGTGCGTATGACTCCGCATCCGGTAGAGTTTGAGCTGTAC
TACAGCGTCTAA

*gyrA* (ECK2223)

ATGAGCGACCTTGCGAGAGAAATTACACCGGTCAACATTGAGGAAGAGCTGA
AGAGCTCCTATCTGGATTATGCGATGTCGGTCATTGTTGGCCGTGCGCTGCC
AGATGTCCGAGATGGCCTGAAGCCGGTACACCGTCGCGTACTTTACGCCAT
GAACGTACTAGGCAATGACTGGAACAAAGCCTATAAAAAATCTGCCCGTGTC
GTTGGTGACGTAATCGGTAAATACCATCCCCATGGTGACTCGGCGGTCTATG
ACACGATCGTCCGCATGGCGCAGCCATTCTCGCTGCGTTATATGCTGGTAGA
CGGTCAGGGTAACTTCGGTTCTATCGACGGCGACTCTGCGGCGGCAATGCG
TTATACGGAAATCCGTCTGGCGAAAATTGCCCATGAACTGATGGCCGATCTC
GAAAAAGAGACGGTCGATTTCGTTGATAACTATGACGGCACGGAAAAAATTC
CGGACGTCATGCCAACCAAAATTCCTAACCTGCTGGTGAACGGTTCTTCCGG
TATCGCCGTAGGTATGGCAACCAACATCCCGCCGCACAACCTGACGGAAGT
CATCAACGGTTGTCTGGCGTATATTGATGATGAAGACATCAGCATTGAAGGG
CTGATGGAACACATCCCGGGGCCGGACTTCCCGACGGCGGCAATCATTAAC
GGTCGTCGCGGTATTGAAGAAGCTTACCGTACCGGTCGCGGCAAGGTGTAT
ATCCGCGCTCGCGCAGAAGTGGAAGTTGACGCCAAAACCGGTCGTGAAACC
ATTATCGTCCACGAAATTCCGTATCAGGTAAACAAAGCGCGCCTGATCGAGA
AGATTGCGGAACTGGTAAAAGAAAAACGCGTGGAAGGCATCAGCGCGCTGC
GTGACGAGTCTGACAAAGACGGTATGCGCATCGTGATTGAAGTGAAACGCG
ATGCGGTCGGTGAAGTTGTGCTCAACAACCTCTACTCCCAGACCCAGTTGCA
GGTTTCTTTCGGTATCAACATGGTGGCATTGCACCATGGTCAGCCGAAGATC
ATGAACCTGAAAGACATCATCGCGGCGTTTGTTCGTCACCGCCGTGAAGTGG
TGACCCGTCGTACTATTTTCGAACTGCGTAAAGCTCGCGATCGTGCTCATAT
CCTTGAAGCATTAGCCGTGGCGCTGGCGAACATCGACCCGATCATCGAACT
GATCCGTCATGCGCCGACGCCTGCAGAAGCGAAAACTGCGCTGGTTGCTAA

TCCGTGGCAGCTGGGCAACGTTGCCGCGATGCTCGAACGTGCTGGCGACGA
TGCTGCGCGTCCGGAATGGCTGGAGCCAGAGTTCGGCGTGCGTGATGGTCT
GTACTACCTGACCGAACAGCAAGCTCAGGCGATTCTGGATCTGCGTTTGCAG
AAACTGACCGGTCTTGAGCACGAAAAACTGCTCGACGAATACAAAGAGCTGC
TGGATCAGATCGCGGAACTGTTGCGTATTCTTGGTAGCGCCGATCGTCTGAT
GGAAGTGATCCGTGAAGAGCTGGAGCTGGTTCGTGAACAGTTCGGTGACAA
ACGTCGTACTGAAATCACCGCCAACAGCGCAGACATCAACCTGGAAGATCTG
ATCACCCAGGAAGATGTGGTCGTGACGCTCTCTCACCAGGGCTACGTTAAGT
ATCAGCCGCTTTCTGAATACGAAGCGCAGCGTCGTGGCGGGAAAGGTAAAT
CTGCCGCACGTATTAAAGAAGACTTTATCGACCGACTGCTGGTGGCGAA
CACTCACGACCATATTCTGTGCTTCTCCAGCCGTGGTCGCGTCTATTCGATG
AAAGTTTATCAGTTGCCGGAAGCCACTCGTGGCGCGCGCGGTCGTCCGATC
GTCAACCTGCTGCCGCTGGAGCAGGACGAACGTATCACTGCGATCCTGCCA
GTGACCGAGTTTGAAGAAGGCGTGAAAGTCTTCATGGCGACCGCTAACGGT
ACCGTGAAGAAAACTGTCCTCACCGAGTTCAACCGTCTGCGTACCGCCGGTA
AAGTGGCGATCAAACTGGTTGACGGCGATGAGCTGATCGGCGTTGACCTGA
CCAGCGGCGAAGACGAAGTAATGC

TGTTCTCCGCTGAAGGTAAAGTGGTGCGCTTTAAAGAGTCTTCTGTCCGTGC
GATGGGCTGCAACACCACCGGTGTTCGCGGTATTCGCTTAGGTGAAGGCGA
TAAAGTCGTCTCTCTGATCGTGCCTCGTGGCGATGGCGCAATCCTCACCGCA
ACGCAAAACGGTTACGGTAAACGTACCGCAGTGGCGGAATACCCAACCAAG
TCGCGTGCGACGAAAGGGGTTATCTCCATCAAGGTTACCGAACGTAACGGTT
TAGTTGTTGGCGCGGTACAGGTAGATGACTGCGACCAGATCATGATGATCAC
CGATGCCGGTACGCTGGTACGTACTCGCGTTTCGGAAATCAGCATCGTGGG
CCGTAACACCCAGGGCGTGATCCTCATCCGTACTGCGGAAGATGAAAACGTA
GTGGGTCTGCAACGTGTTGCTGAACCGGTTGACGAGGAAGATCTGGATACC
ATCGACGGCAGTGCCGCGGAAGGGGACGATGAAATCGCTCCGGAAGTGGA
CGTTGACGACGAGCCAGAAGAAGAATAA

---

[a.] The protospacer sequences are highlighted in blue and the PAMs are highlighted in red.
[b.] The protospacer of the second gRNA (targeting *gyrA*) is highlighted with a single underline, and its corresponding PAM is marked with a double underline.

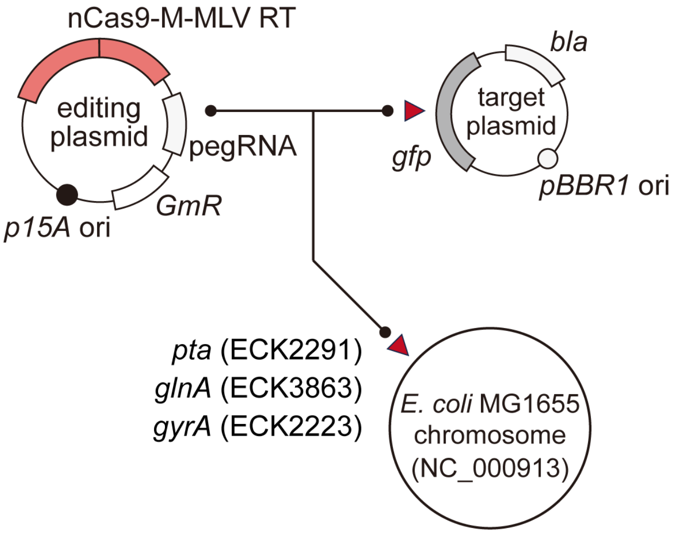


**Supplementary Fig. 1. Experimental design for assessing prime editing capacity.** Customized PE2 was used to target either a plasmid-carrying *gfp* gene or chromosomal loci, including *pta*, *glnA* and *gyrA*. The PE2 effector comprises a fusion of *Streptococcus pyogenes* nCas9 and Moloney murine leukemia virus RT, together with a pegRNA containing the designed edits.

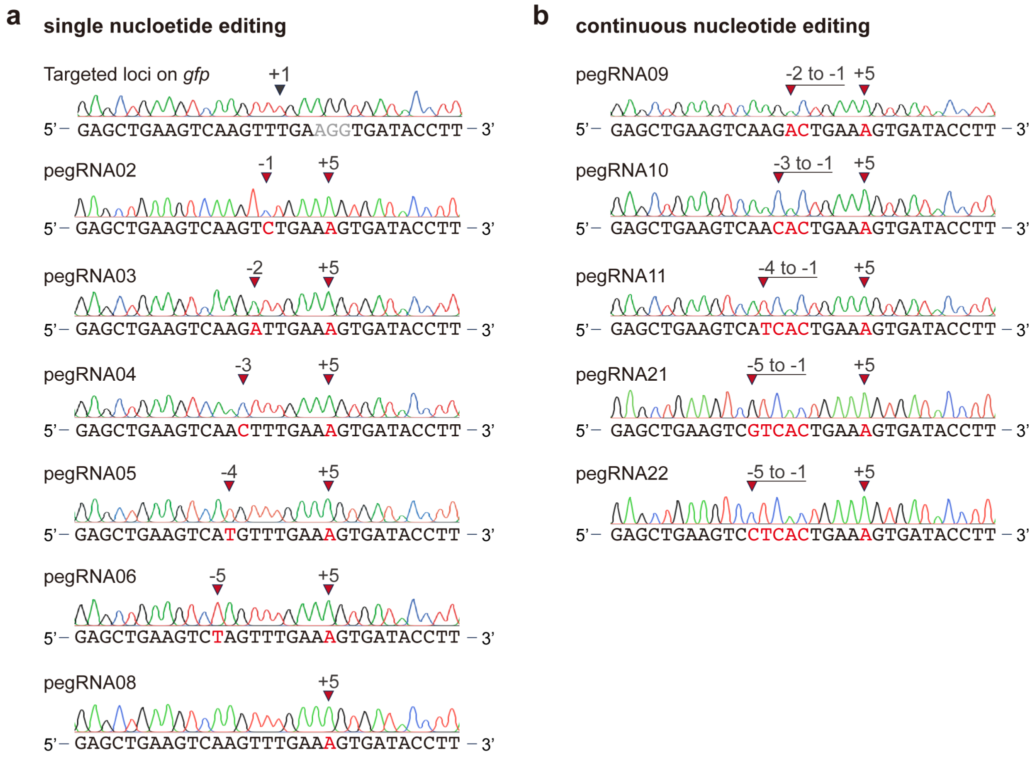


**Supplementary Fig. 2. Additional Sanger sequencing results of prime edited *gfp*. (a)** Sequencing results of single-nucleotide substitution with pegRNAs (pegRNA 02 to pegRNA06 and pegRNA08) and **(b)** continuous nucleotide substitutions (pegRNA09 to pegRNA11, pegRNA21 and pegRNA22) containing 13-nt PBS. The edited positions were indicated by the arrows, and edited bases were highlighted in red.

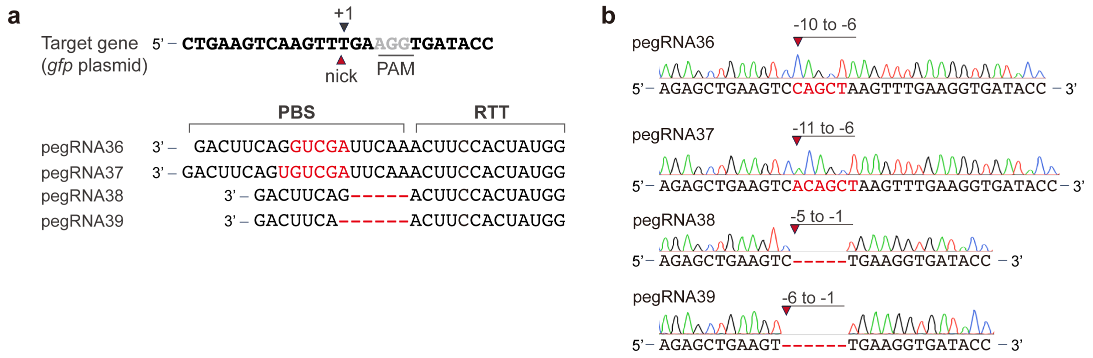


**Supplementary Fig. 3. Additional Sanger sequencing results of continuous 5- and 6-nt insertions and deletions at *gfp*. (a)** Design of pegRNAs with 5-nt insertion (pegRNA36), 6-nt insertion (pegRNA37), 5-nt deletion (pegRNA38) and 6-nt deletion (pegRNA39). **(b)** Sanger sequencing results. The edited positions were indicated by the arrows, and edited bases were highlighted in red.

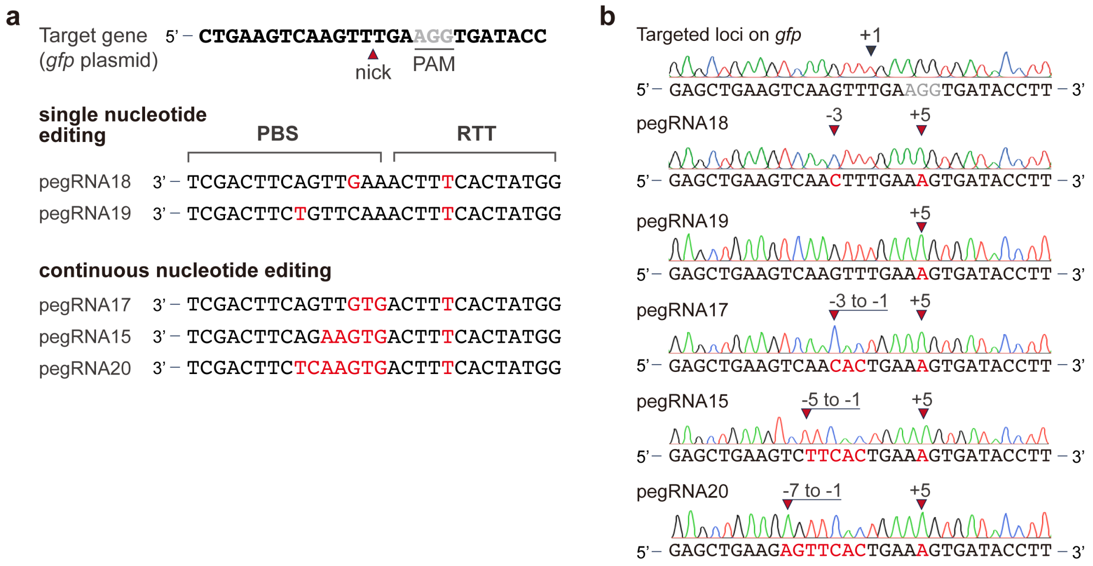


**Supplementary Fig. 4. Prime editing with 15-nt PBS pegRNAs. (a)** Design of pegRNAs with single-nucleotide substitution (pegRNA18 and pegRNA19) or continuous substitutions (pegRNA15, pegRNA17 and pegRNA20) with 15-nt PBS. **(b)** Sanger sequencing results. The edited positions were indicated by the arrows, and edited bases were highlighted in red.

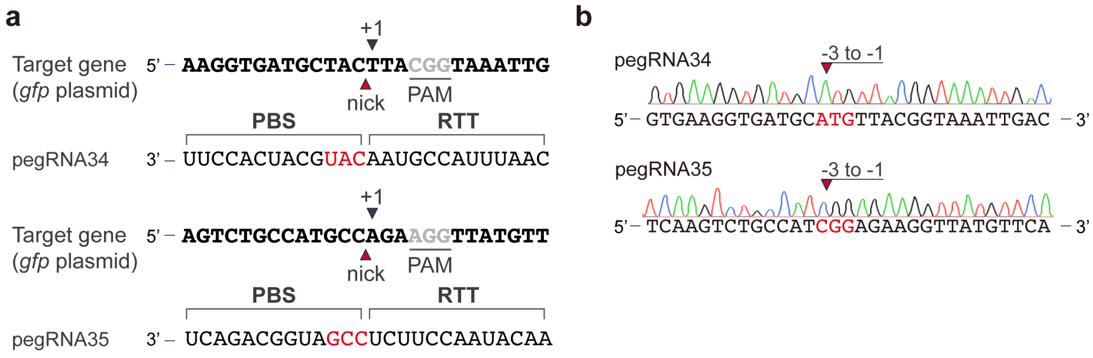


**Supplementary Fig. 5. Sanger sequencing results of 3-nt continuous substitutions at two additional *gfp* sites. (a)** Design of pegRNA34 and pegRNA35. **(b)** Sanger sequencing results. The edited positions were indicated by the arrows, and edited bases were highlighted in red.

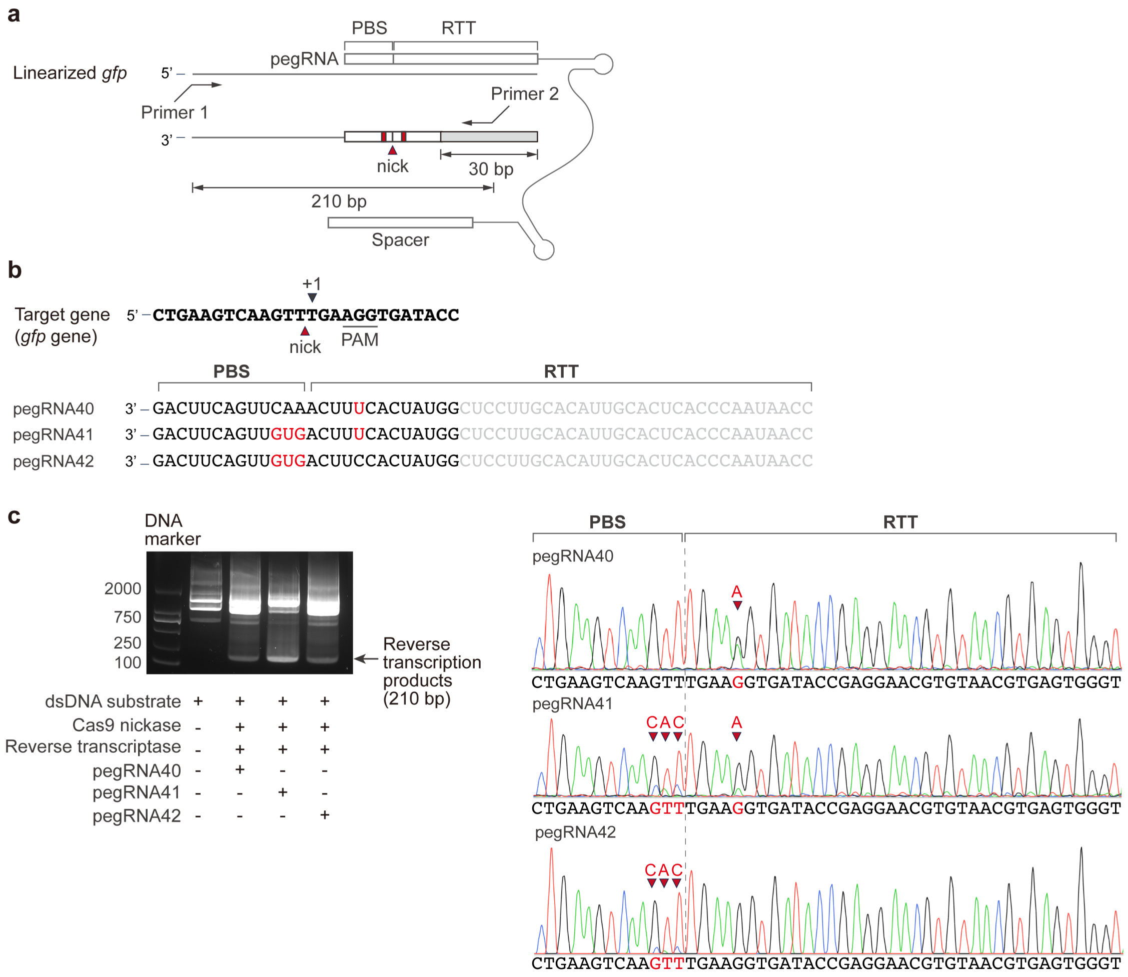


**Supplementary Fig. 6. *In vitro* validation of non-canonical reverse transcription pathway using purified components. (a)** Schematic illustration of the *in vitro* test using a linearized *gfp* DNA substrate. Primer 1 (XIA-PRC-554) and primer 2 (XIA-PRC-555) were used to amplify the target region, with primer 1 annealing to the target DNA, while primer 2 is designed to anneal only to the reverse transcription products. **(b)** pegRNA designs. pegRNA40 was designed for +5 substitution, while pegRNA41 and pegRNA42 were designed for upstream 3-nt continuous substitutions with or without +5 substitution. A prolonged RTT is designed with a 30 bp random sequence for subsequent PCR of the reverse transcription product. **(c)** Agarose gel electrophoresis of amplified products and Sanger sequencing results. The sequencing results show reverse transcription products containing editing in RTT (pegRNA40), in PBS (pegRNA42), and both (pegRNA41). The edited positions were indicated by the arrows, and edited bases were highlighted in red.

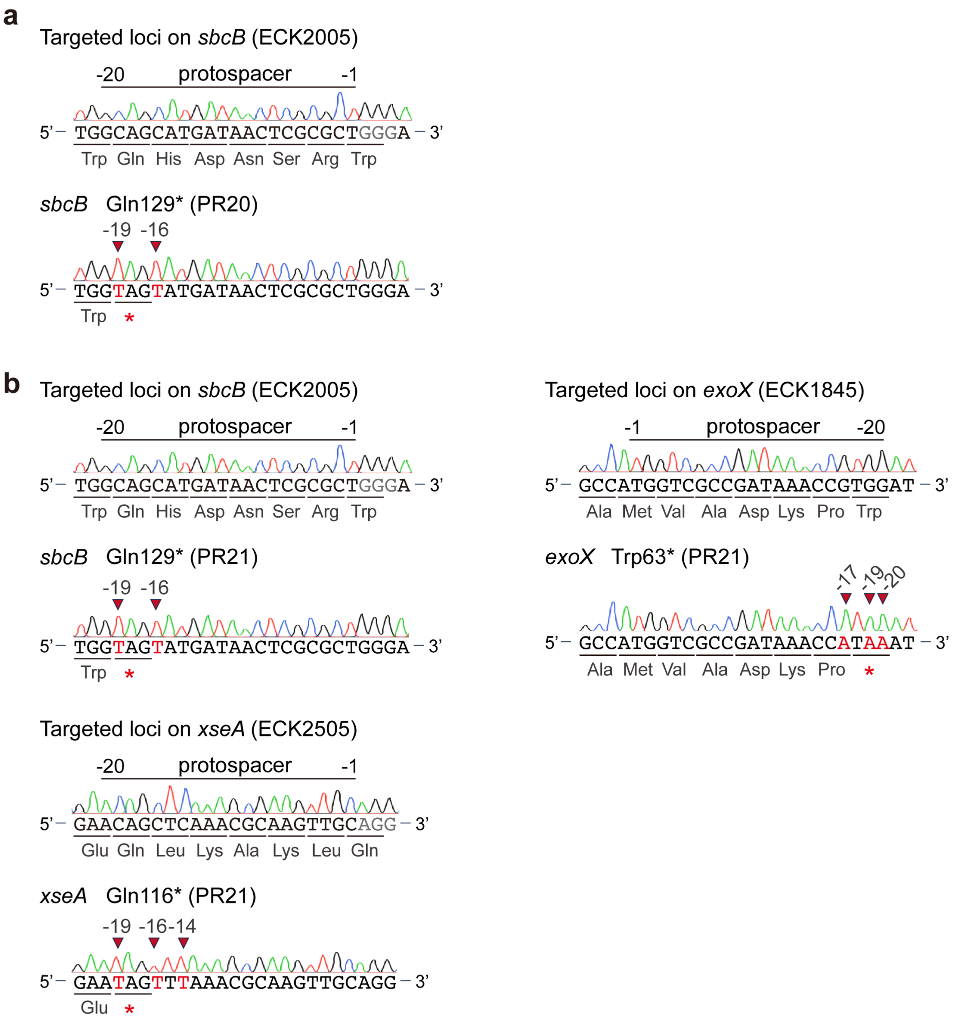


**Supplementary Fig. 7. Disruption of DNA exonucleases with base editing. (a)** Sequencing results of edited *sbcB* in PR20, and **(b)** simultaneous editing of *sbcB*, *xseA* and *exoX* in PR21. The edited positions were indicated by the arrows, and the altered amino acids and edited bases were marked in red.

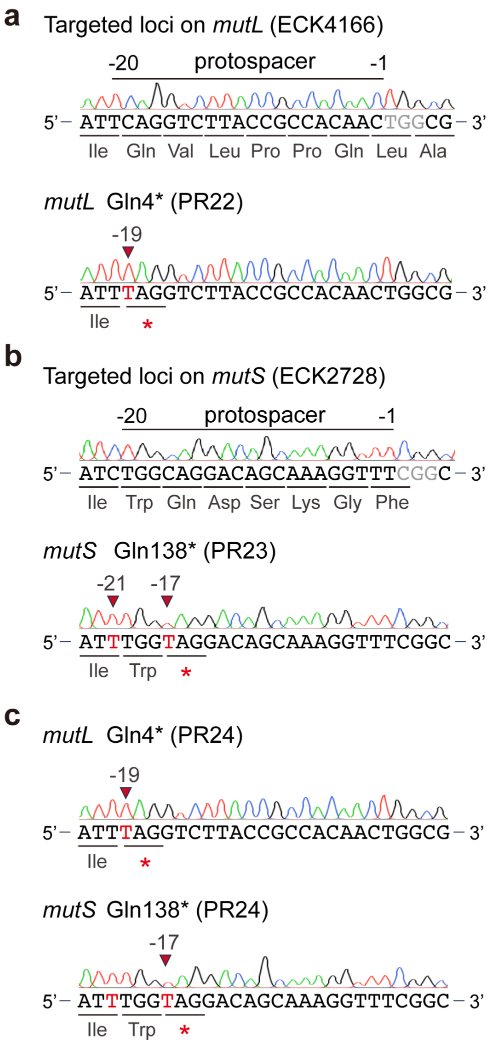


**Supplementary Fig. 8. Disruption of MMR. (a)** Sequencing results of edited *mutL* in PR22, **(b)** *mutS* in PR23, and **(c)** both genes in PR24. The edited positions were indicated by the arrows, and the altered amino acids and edited bases were marked in red.

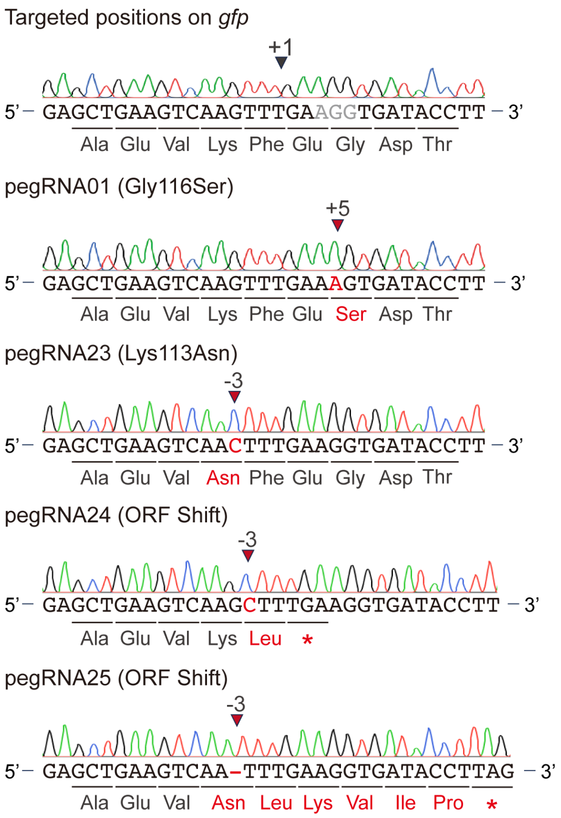


**Supplementary Fig. 9. Sanger sequencing validation of prime edited *gfp* with pegRNA 01 and pegRNA23 to pegRNA25.** Sequencing results confirming the intended mutation introduced by prime editing were used for fluorescence assays. The edited positions were indicated by the arrows, and the altered amino acids and edited bases were marked in red.

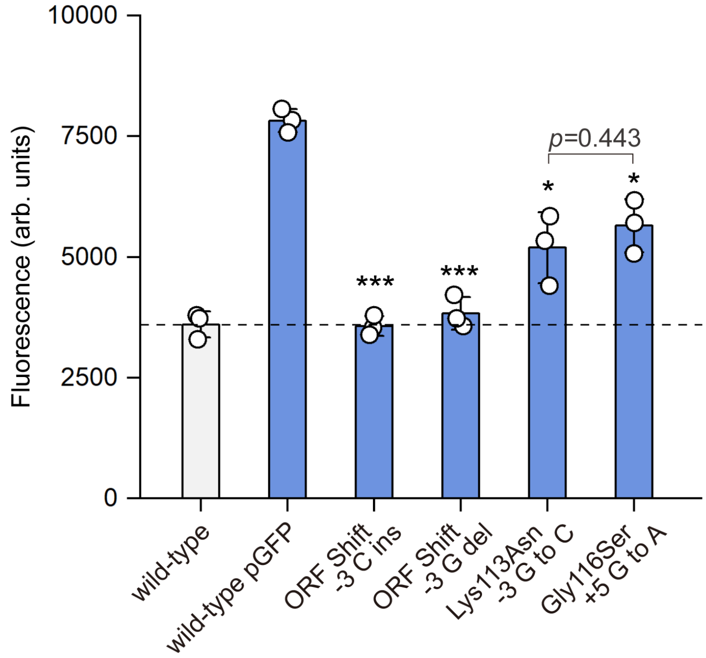


**Supplementary Fig. 10. Fluorescence analysis of edited *gfp* strains.** Fluorescence intensities of *E. coli* strains expressing wild-type *gfp* (wild-type pGFP) or edited *gfp*. The wild-type strain without a plasmid served as a negative control. Data are presented as mean values ± SD (n = 3 biological replicates). Statistical significance is determined by comparing each variant to wild-type pGFP, *t*-test (*, $p<0.05$; ***, $p<0.001$, unpaired and two-sided), and the exact *p*-values are provided in Source Data. Source data are provided as a Source Data file.

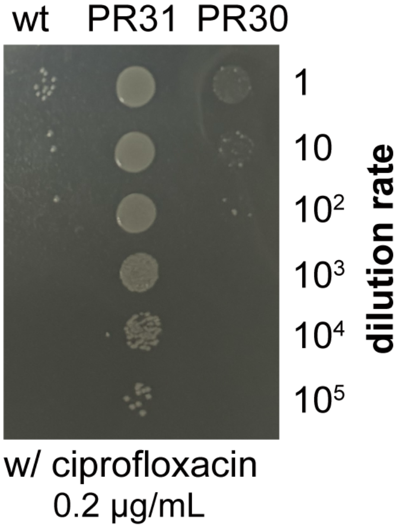


**Supplementary Fig. 11. Ciprofloxacin susceptibility test of *gyrA* mutant strains 0.2 µg/mL ciprofloxacin.** Strains were adjusted to $OD_{600}$=1, serially diluted from $10^{-1}$ to $10^{-5}$, and spotted onto LB plates containing 0.2 µg/mL ciprofloxacin. The wild-type (wt) strain served as a control.
